## Supplementary Figures for "Multiple protein-DNA interfaces unravelled by evolutionary information, physico-chemical and geometrical properties"

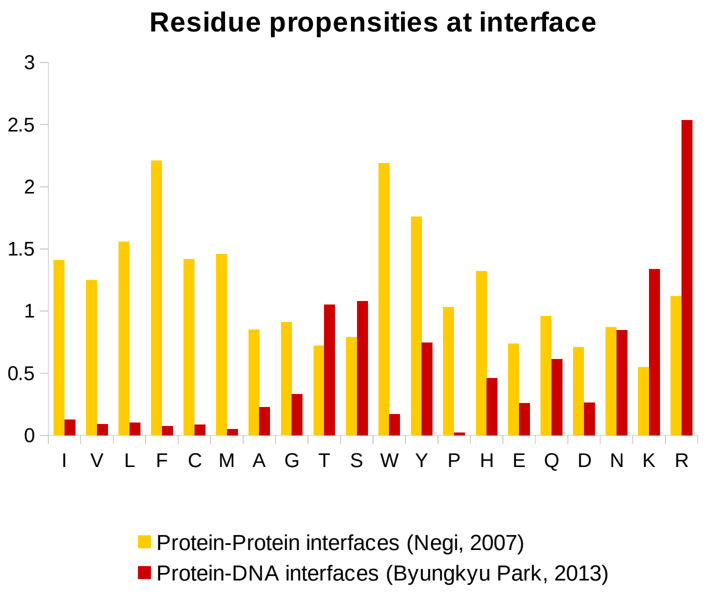


Figure S1: **Interface propensities.** The protein-DNA (in red, [**?**]) and protein-protein (in orange, [**?**]) scales are shown on the same plot for visual comparison. Amino acids are ordered from the most to the least hydrophobic one, based on the Kyte and Doolittle hydrophobicity scale [**?**].

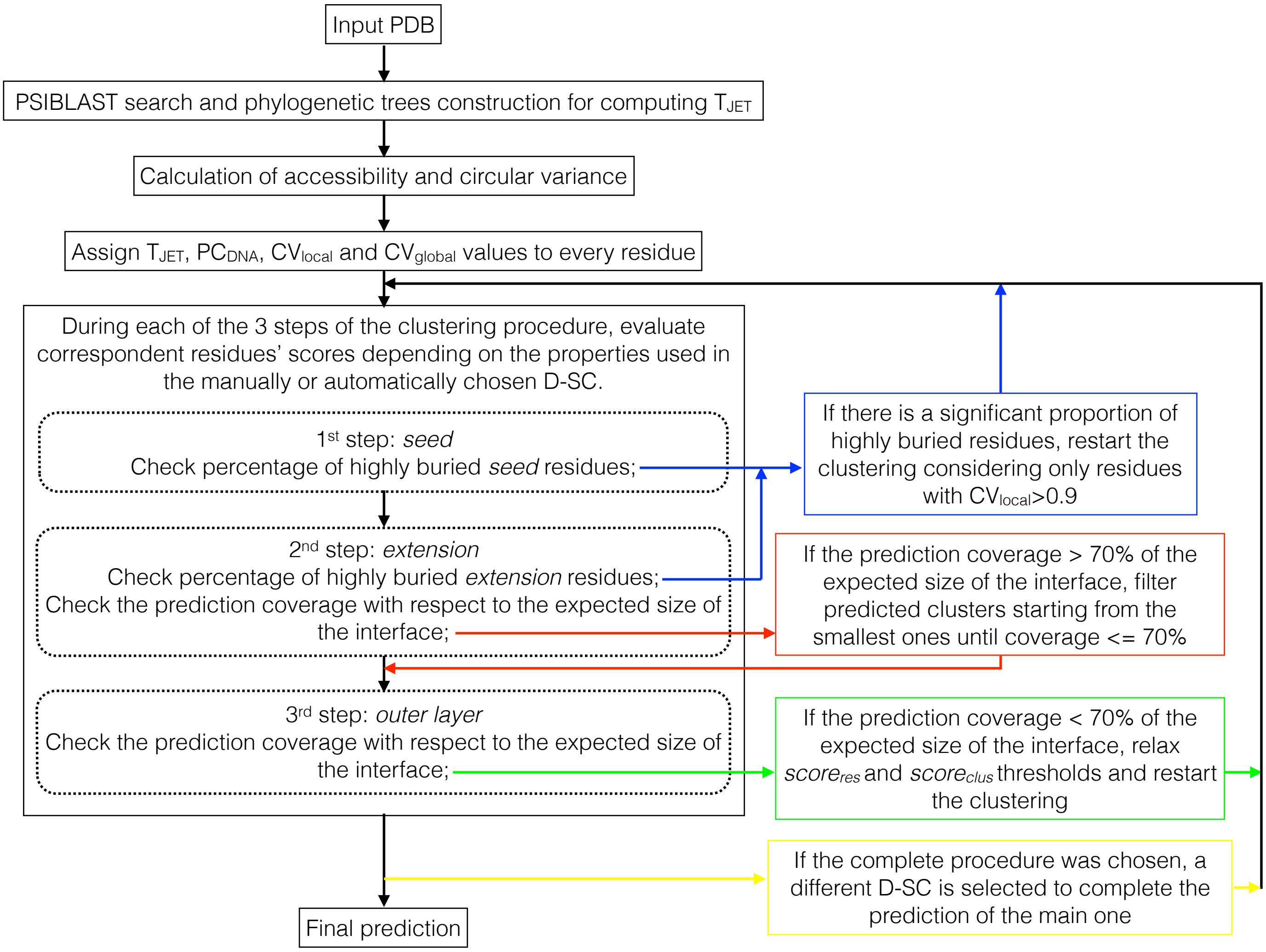


Figure S2: **JET$^2_{\mathrm{DNA}}$ pipeline.** General JET$^2_{\mathrm{DNA}}$ pipeline, whatever the scoring scheme chosen. In black, the mandatory steps. In blue, procedure to avoid buried small ligand binding pockets. In red, filtering of the putative false positive clusters. In green, relaxing of the thresholds for too small predicted clusters, which do not respect the expecting size of the interface. In yellow, the possibility to complete the main prediction with a second one obtained by a different scoring scheme.

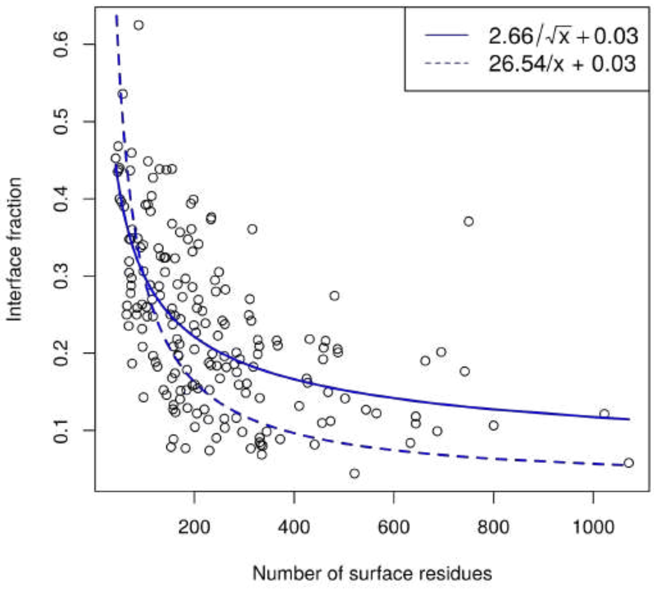


Figure S3: **Expected size of the interface**. Plot of the $f_{intfrac}$ function relating surface size and fraction of the surface covered by the interface. Each circle is the percentage of interface residues versus the total number of surface residues for a given protein comprised in the HR-PDNA187 dataset. The solid line corresponds to the function that best approximates the circles distribution. The dotted line corresponds to the function that best approximates percentage of interface residues versus the total number of surface residues for protein-protein interfaces [**?**] based on a dataset of 1256 protein chains [**?**].

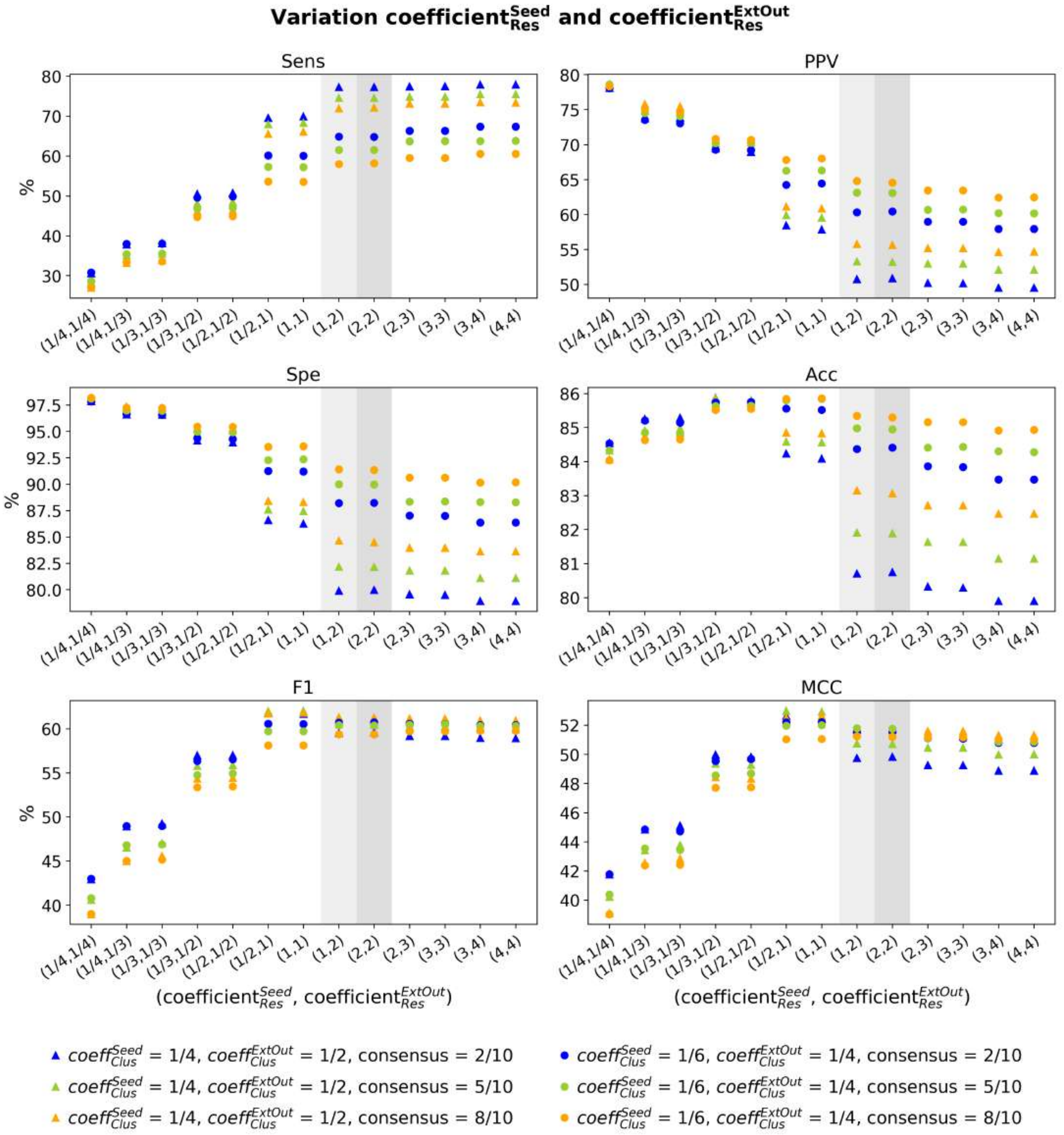


Figure S4: **Influence of the two parameters coefficient$_{Res}^{Seed}$ and coefficient$_{Res}^{ExtOut}$ on the predictive performance of JET$^2_{\mathrm{DNA}}$.** These coefficients are multiplied by $f_{intfrac}$ to determine the two residue thresholds score$_{Res}^{Seed}$ and score$_{Res}^{ExtOut}$. Namely, only residues showing a score $\geq$ score$_{Res}^{Seed}$ (resp. score$_{Res}^{ExtOut}$) are considered by the JET$^2_{\mathrm{DNA}}$ algorithm during the detection of the seed (resp. extension). Two different sets of coefficient$_{Clus}^{Seed}$, coefficient$_{Clus}^{ExtOut}$ and Confidence$_{ClusterSize}$ were tried (triangles and circles) to test the stability of the results with respect to the other thresholds used in the algorithm. JET$^2_{\mathrm{DNA}}$ predictions obtained with a consensus of 2, 5 and 8 runs out of 10 are reported in blue, green and orange respectively. The calculation was performed on the HR-PDNA187 dataset. Default values chosen in JET$^2_{\mathrm{DNA}}$ for the first stage (not-relaxed values) and the second stage (relaxed values) of prediction are highlighted in lightgrey and grey, respectively.

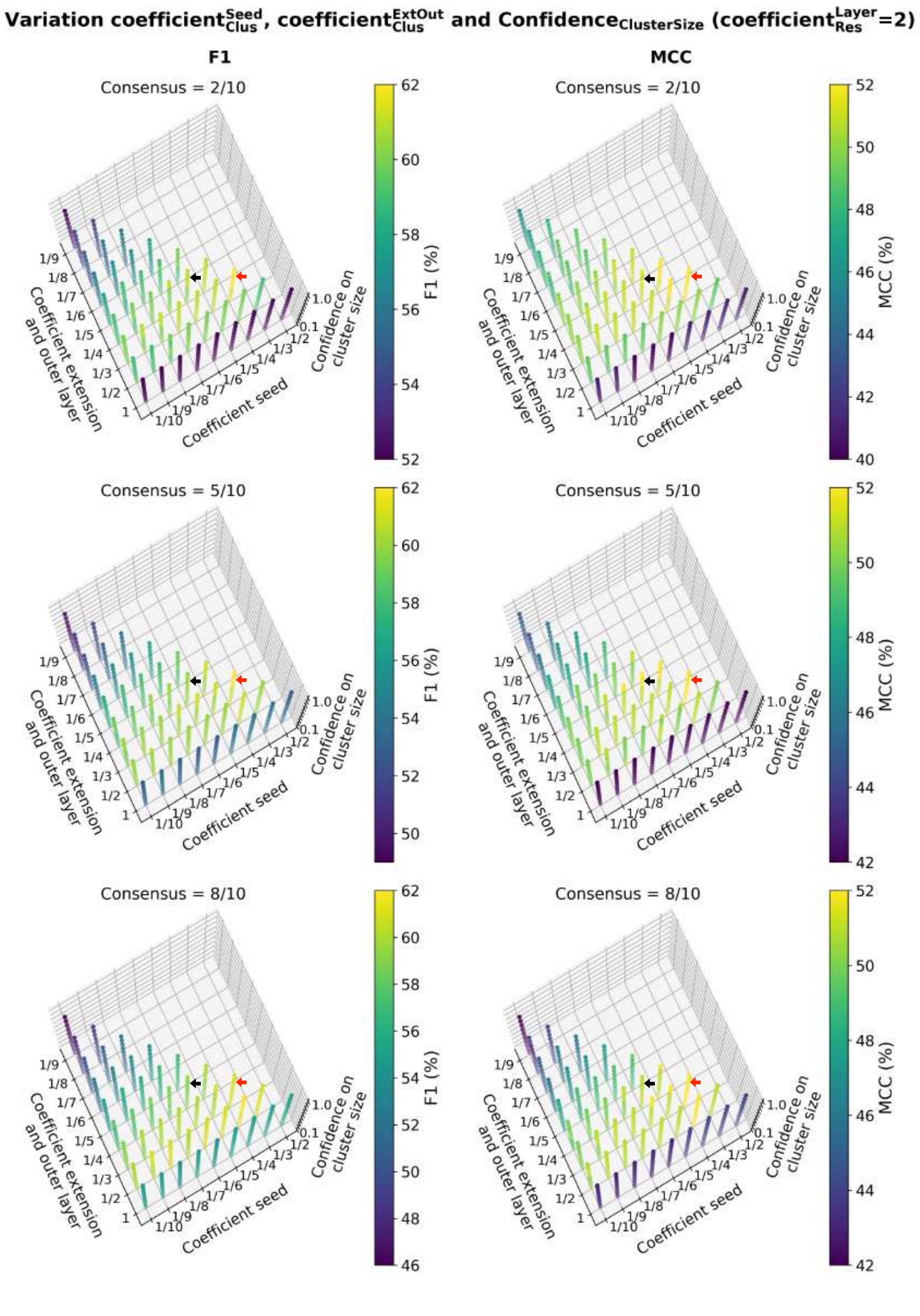


Figure S5: **Influence of the three parameters coefficient$^{Seed}_{Clus}$, coefficient$^{ExtOut}_{Clus}$ and confidence$_{clusSize}$ on MCC and F1 values.** coefficient$^{Seed}_{Clus}$ and coefficient$^{ExtOut}_{Clus}$ are multiplied by $f_{intfrac}$ to determine the two cluster thresholds score$^{Seed}_{Clus}$ and score$^{ExtOut}_{Clus}$. Namely, a given seed (resp. extension) is grown until the averaged score of the predicted cluster reaches down score$^{Seed}_{Clus}$ (resp. score$^{ExtOut}_{Clus}$). The third threshold represents the confidence on the cluster size (confidence$_{clusSize}$). Clusters with a size below this confidence level are filtered out. The statistical value is coded by the color. Results obtained from a consensus of 2, 5 and 8 runs out of 10 are reported on top, at the center and bottom, respectively. The calculation was performed on the HR-PDNA187 dataset. Default values chosen in JET$^2_{\text{DNA}}$ for the first stage (not-relaxed values) and the second stage (relaxed values) of prediction are highlighted in black and red, respectively.

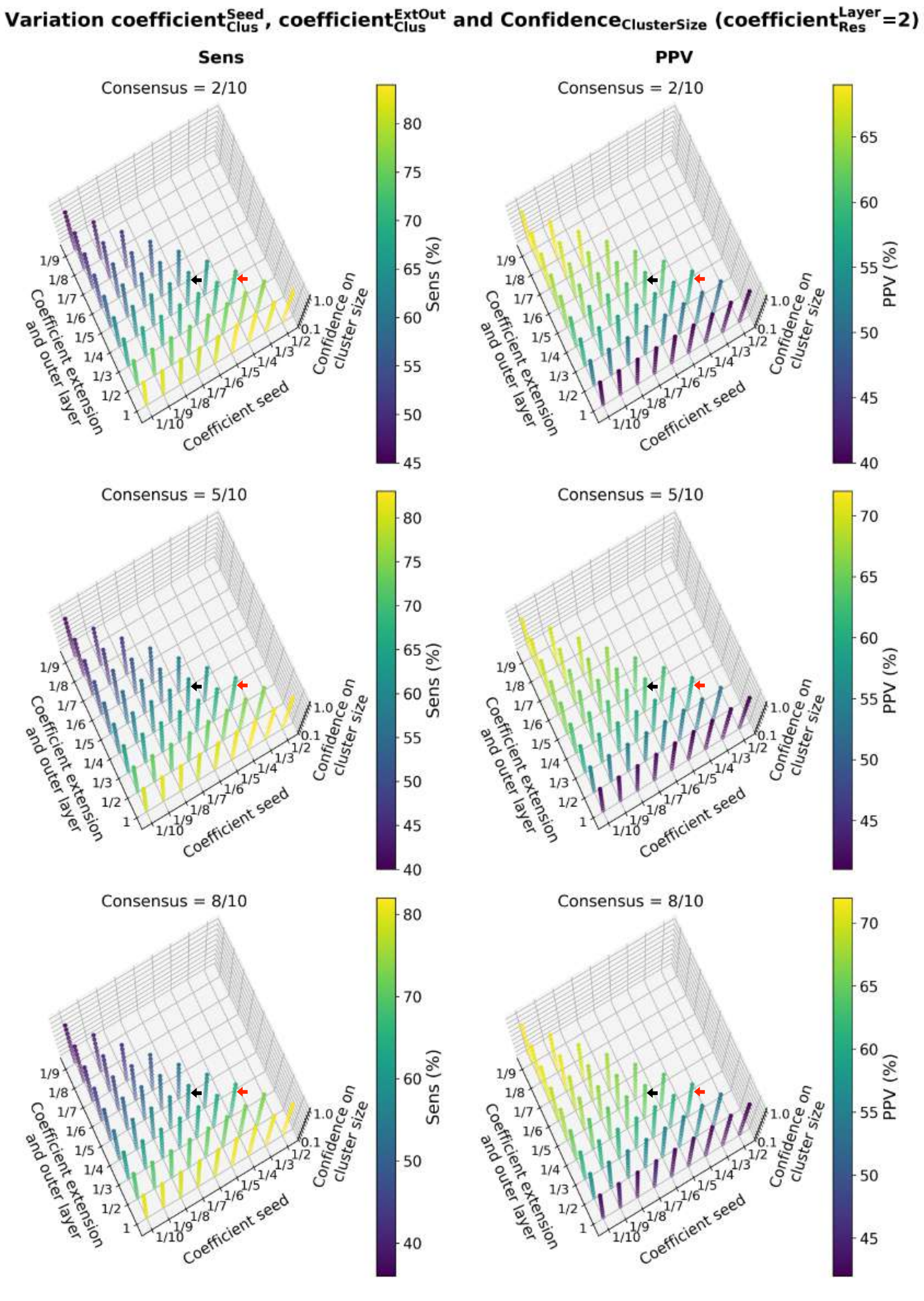


Figure S6: **Influence of the three parameters coefficient$^{Seed}_{Clus}$, coefficient$^{ExtOut}_{Clus}$ and confidence$_{clusSize}$ on the sensitivity and PPV values.** coefficient$^{Seed}_{Clus}$ and coefficient$^{ExtOut}_{Clus}$ are multiplied by $f_{intfrac}$ to determine the two cluster thresholds score$^{Seed}_{Clus}$ and score$^{ExtOut}_{Clus}$. Namely, a given seed (resp. extension) is grown until the averaged score of the predicted cluster reaches down score$^{Seed}_{Clus}$ (resp. score$^{ExtOut}_{Clus}$). The third threshold represents the confidence on the cluster size (confidence$_{clusSize}$). Clusters with a size below this confidence level are filtered out. The statistical value is coded by the color. Results obtained from a consensus of 2, 5 and 8 runs out of 10 are reported on top, at the center and bottom, respectively. The calculation was performed on the HR-PDNA187 dataset. Default values chosen in JET$^2_{\mathrm{DNA}}$ for the first stage (not-relaxed values) and the second stage (relaxed values) of prediction are highlighted in black and red, respectively.

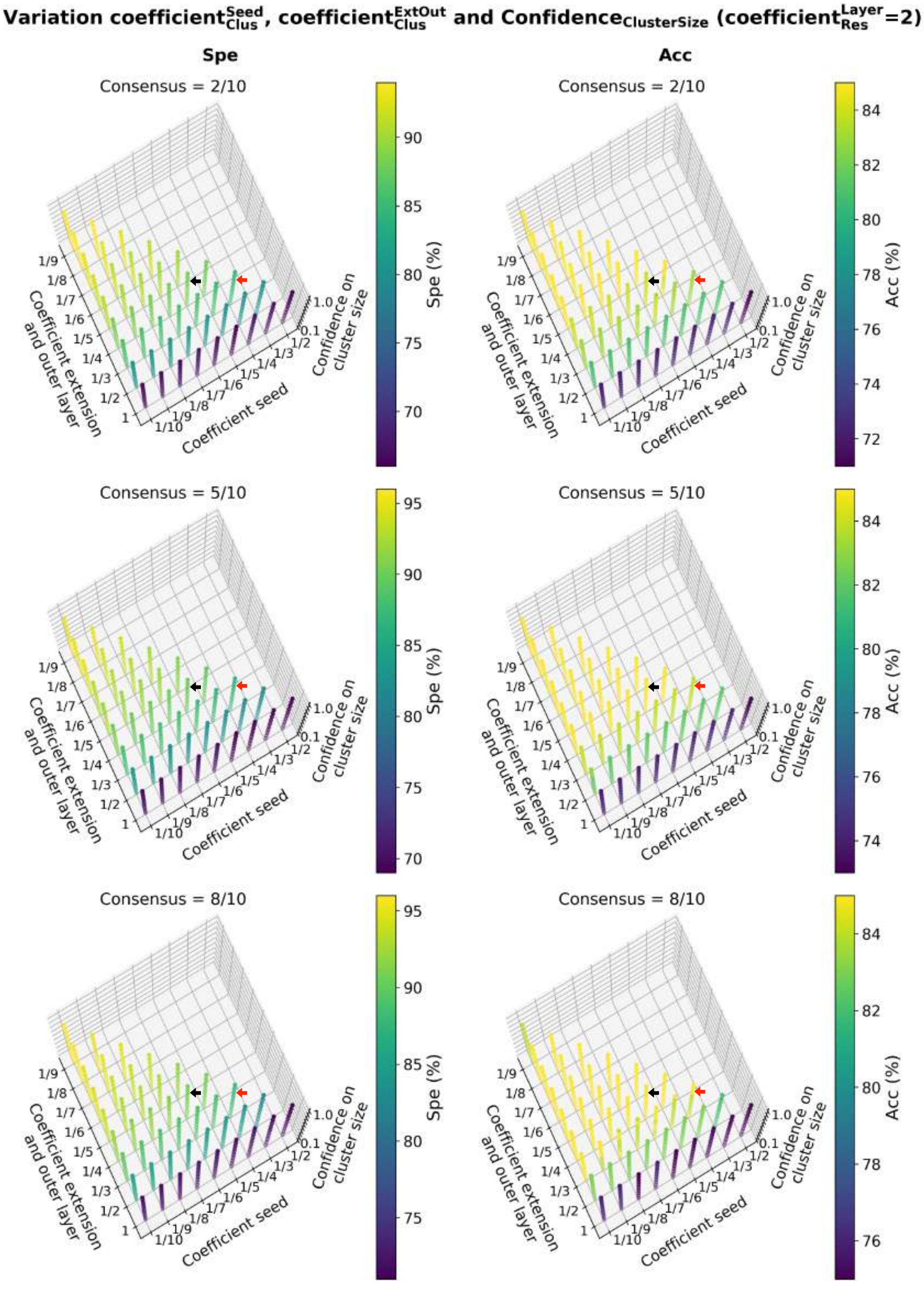


Figure S7: **Influence of the three parameters coefficient$^{Seed}_{Clus}$, coefficient$^{ExtOut}_{Clus}$ and confidence$_{clusSize}$ on specificity and accuracy values.** coefficient$^{Seed}_{Clus}$ and coefficient$^{ExtOut}_{Clus}$ are multiplied by $f_{intfrac}$ to determine the two cluster thresholds score$^{Seed}_{Clus}$ and score$^{ExtOut}_{Clus}$. Namely, a given seed (resp. extension) is grown until the averaged score of the predicted cluster reaches down score$^{Seed}_{Clus}$ (resp. score$^{ExtOut}_{Clus}$). The third threshold represents the confidence on the cluster size (confidence$_{clusSize}$). Clusters with a size below this confidence level are filtered out. The statistical value is coded by the color. Results obtained from a consensus of 2, 5 and 8 runs out of 10 are reported on top, at the center and bottom, respectively. The calculation was performed on the HR-PDNA187 dataset. Default values chosen in JET$^2_{\mathrm{DNA}}$ for the first stage (not-relaxed values) and the second stage (relaxed values) of prediction are highlighted in black and red, respectively.

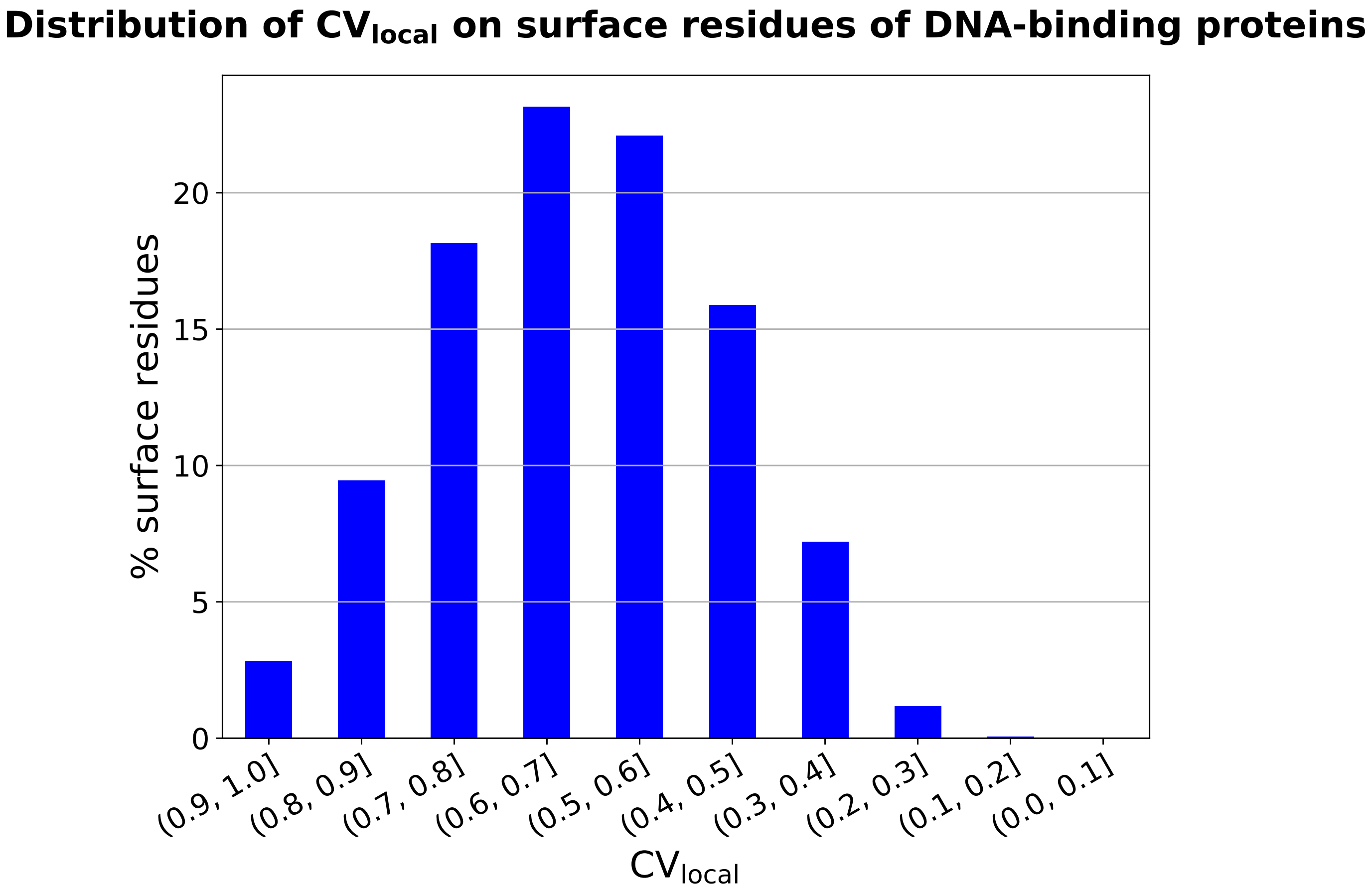


Figure S8: **Distribution of $CV_{local}$ values on DNA-binding protein surfaces.** Histogram of the percentage of surface residues having a $CV_{local}$ value comprised in one of the ten intervals between [0, 1]. The calculation was performed on the HR-PDNA187 dataset. Surface residues are defined as the ones showing a relative solvent accessibility $\geq 5\%$. High values of $CV_{local}$ indicate locally buried residues.

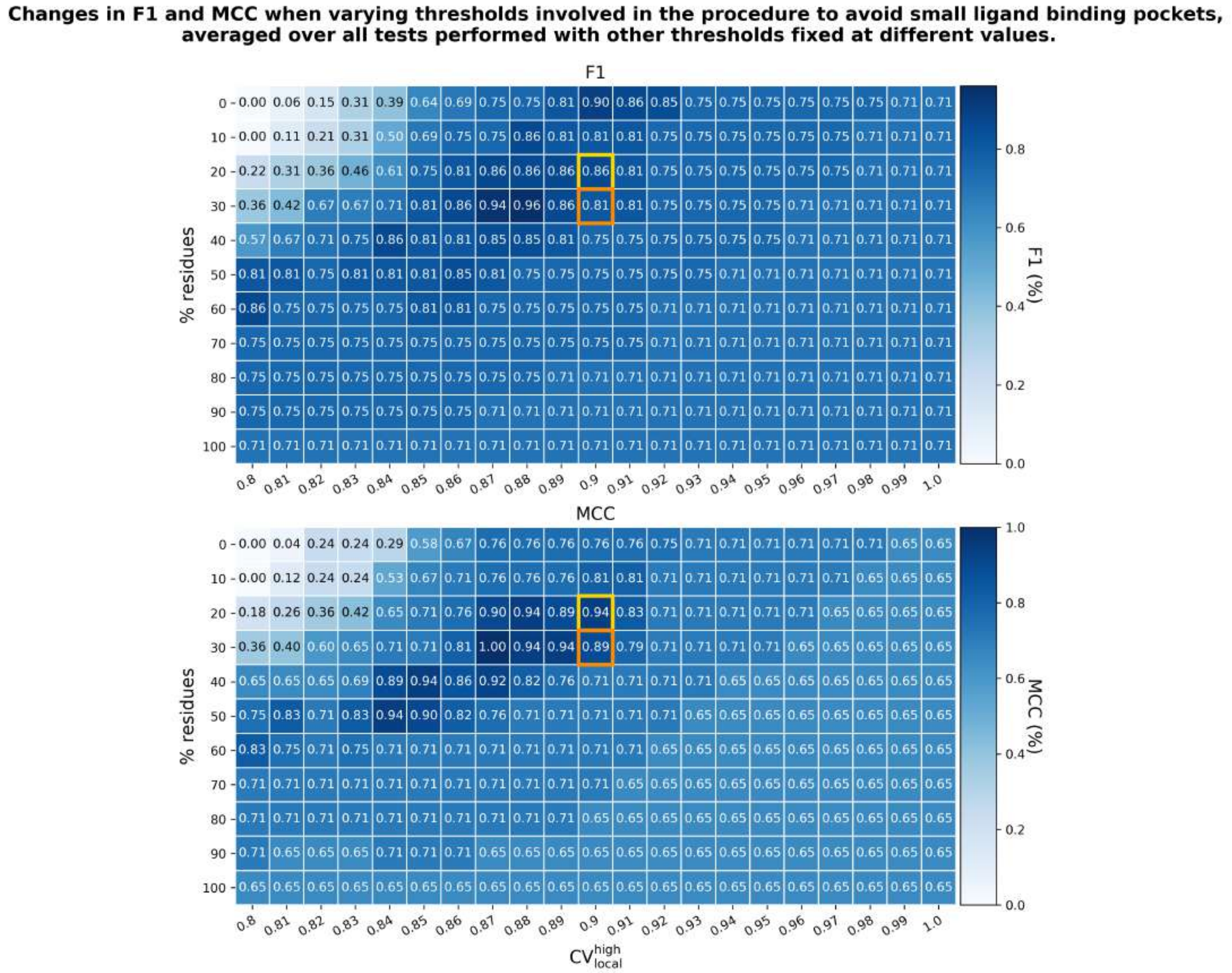


Figure S9: **Influence of the two parameters involved in the procedure to avoid the prediction of small ligand binding pockets on F1 and MCC values.** The $CV_{local}{}^{high}$ (x-axis) and the percentage of residues allowed to show $CV_{local} \geq CV_{local}{}^{high}$ (y-axis) are the two parameters involved in the procedure to avoid the prediction of small ligand binding pockets. In each cell is reported the F1 (on top) or the MCC value (at the bottom) averaged over the six 2D plots in Supp. Fig. 10-11. F1 and MCC values in each 2D plot in Supp. Fig. 10-11 were first normalized with respect to the minimum and maximum values over the plot. The calculation was performed on the HR-PDNA187 dataset. Default values chosen in $JET^2_{DNA}$ for D-SC1 and D-SC2 are highlighted in yellow and orange, respectively.

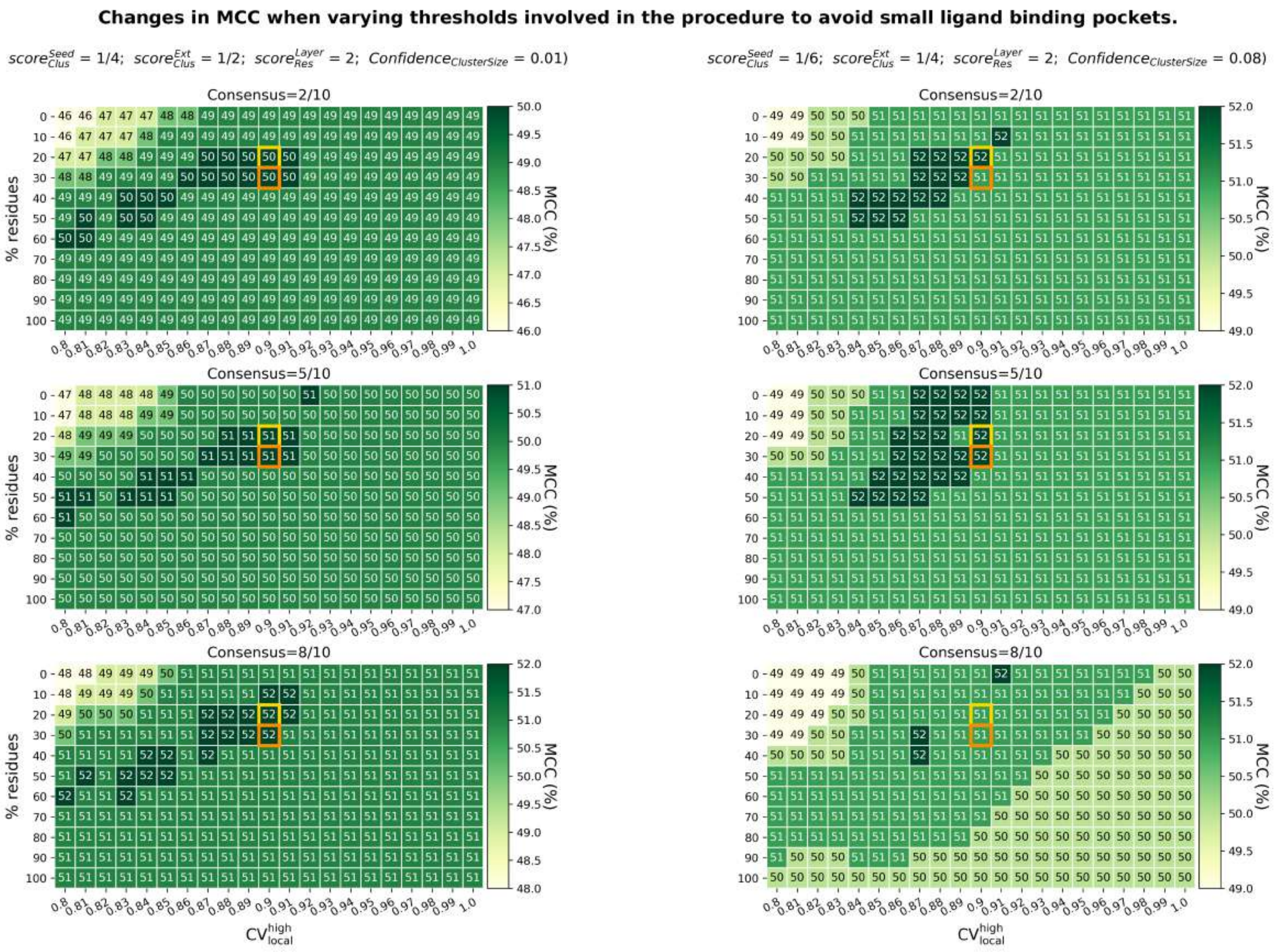


Figure S10: **Influence of the two parameters involved in the procedure to avoid the prediction of small ligand binding pockets on MCC values.** The $\mathrm{CV_{local}}^{\mathrm{high}}$ (x-axis) and the percentage of residues allowed to show $\mathrm{CV_{local}} \geq \mathrm{CV_{local}}^{\mathrm{high}}$ (y-axis) are the two parameters involved in the procedure to avoid the prediction of small ligand binding pockets. In each cell is reported the averaged MCC value computed on the HR-PDNA187 dataset. Two different sets of $\mathrm{score}_{Clus}^{Seed}$, $\mathrm{score}_{Clus}^{ExtOut}$, $\mathrm{score}_{Res}^{Layer=seed,extension,outer\ layer}$ and $\mathrm{Confidence}_{ClusterSize}$ were tried, reported on the left and right columns respectively, to test the stability of results with respect to these other thresholds used in the algorithm. In each column, three 2D plots are reported for predictions obtained with a consensus of 2, 5 and 8 runs out of 10, respectively. Default values chosen in $\mathrm{JET}^2_{\mathrm{DNA}}$ for D-SC1 and D-SC2 are highlighted in yellow and orange, respectively.

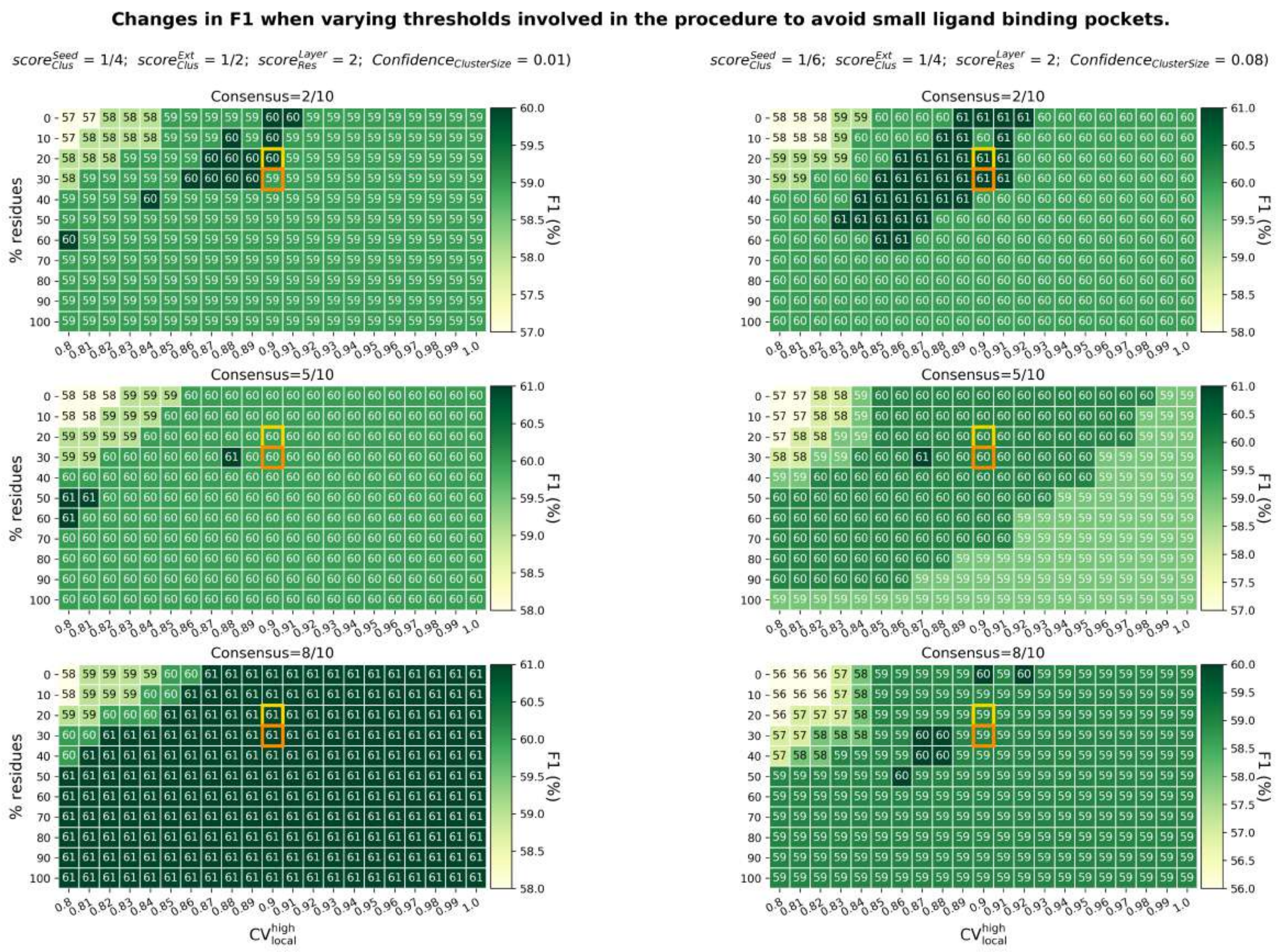


Figure S11: **Influence of the two parameters involved in the procedure to avoid the prediction of small ligand binding pockets on F1 values.** The $\mathrm{CV_{local}}^{\mathrm{high}}$ (x-axis) and the percentage of residues allowed to show $\mathrm{CV_{local}} \geq \mathrm{CV_{local}}^{\mathrm{high}}$ (y-axis) are the two parameters involved in the procedure to avoid the prediction of small ligand binding pockets. In each cell is reported the averaged F1 value computed on the HR-PDNA187 dataset. Two different sets of $\mathrm{score}^{Seed}_{Clus}$, $\mathrm{score}^{ExtOut}_{Clus}$, $\mathrm{score}^{Layer=seed,extension,outer\ layer}_{Res}$ and $\mathrm{Confidence}_{ClusterSize}$ were tried, reported on the left and right columns respectively, to test the stability of results with respect to these other thresholds used in the algorithm. In each column, three 2D plots are reported for predictions obtained with a consensus of 2, 5 and 8 runs out of 10, respectively. Default values chosen in $\mathrm{JET}^2_{\mathrm{DNA}}$ for D-SC1 and D-SC2 are highlighted in yellow and orange, respectively.

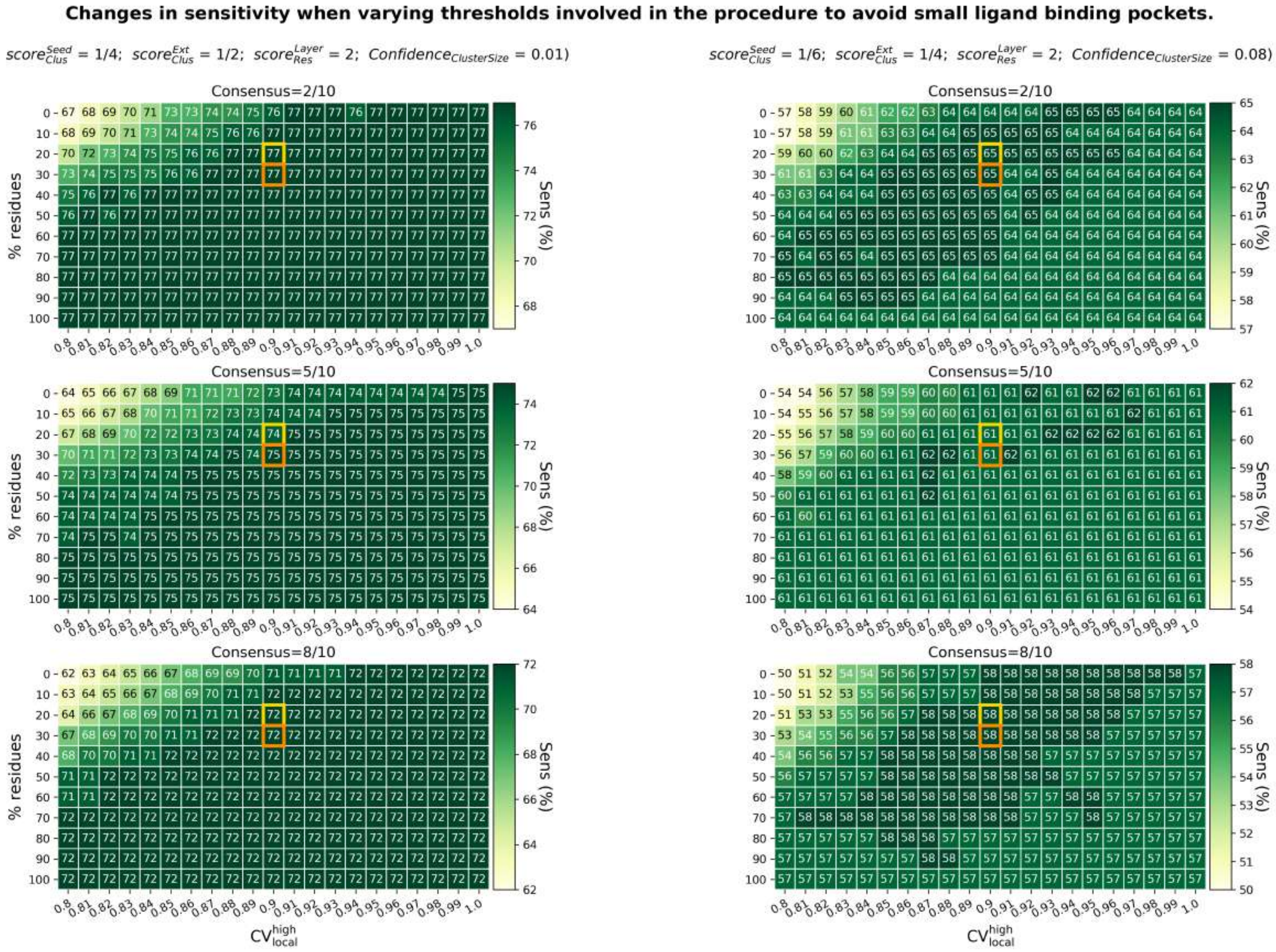


Figure S12: **Influence of the two parameters involved in the procedure to avoid the prediction of small ligand binding pockets on sensitivity values.** The $CV_{local}{}^{high}$ (x-axis) and the percentage of residues allowed to show $CV_{local} \geq CV_{local}{}^{high}$ (y-axis) are the two parameters involved in the procedure to avoid the prediction of small ligand binding pockets. In each cell is reported the averaged sensitivity value computed on the HR-PDNA187 dataset. Two different sets of $score_{Clus}^{Seed}$, $score_{Clus}^{ExtOut}$, $score_{Res}^{Layer=seed,extension,outer\ layer}$ and $Confidence_{ClusterSize}$ were tried, reported on the left and right columns respectively, to test the stability of results with respect to these other thresholds used in the algorithm. In each column, three 2D plots are reported for predictions obtained with a consensus of 2, 5 and 8 runs out of 10, respectively. Default values chosen in $JET^2_{DNA}$ for D-SC1 and D-SC2 are highlighted in yellow and orange, respectively.

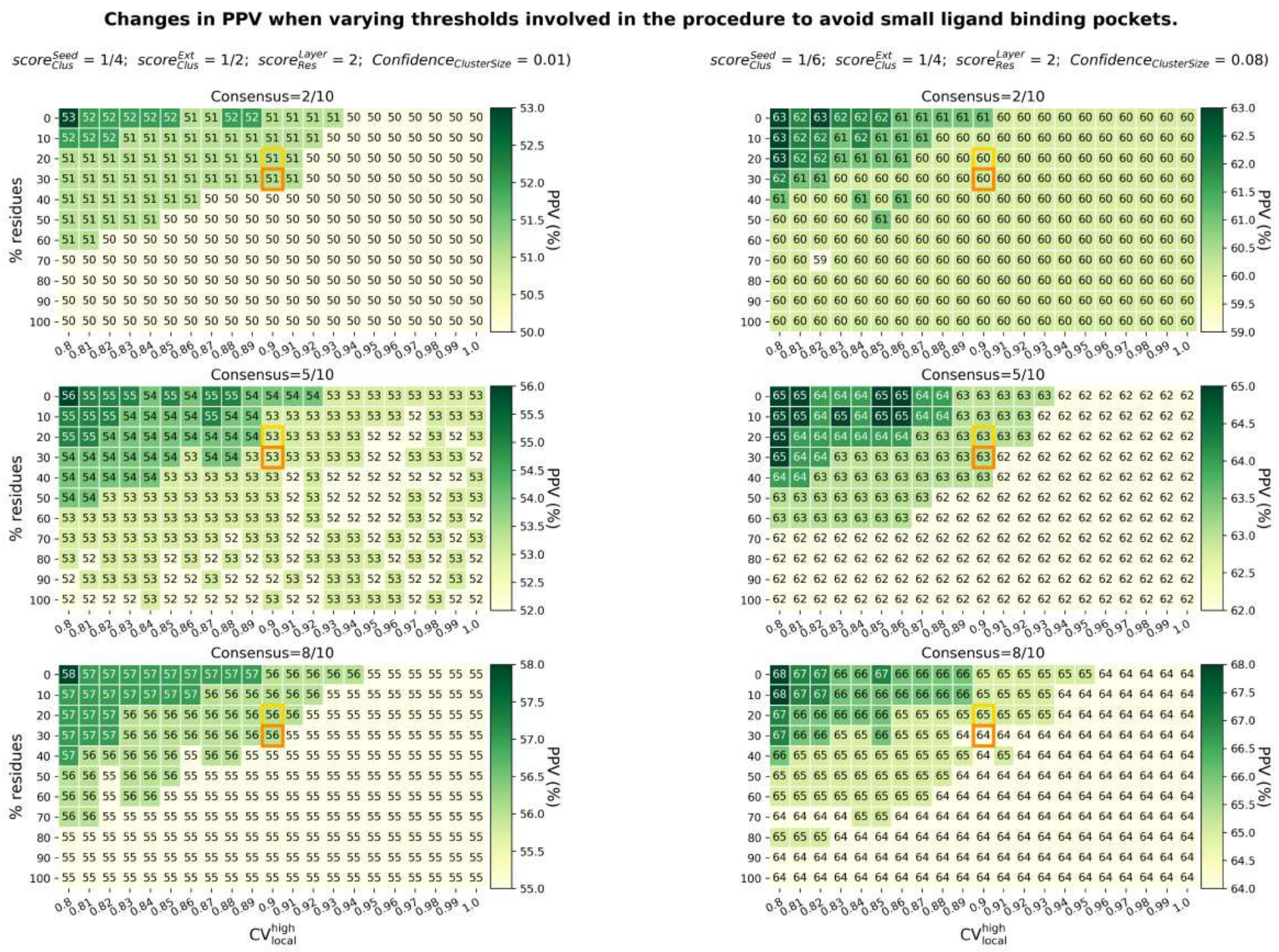


Figure S13: **Influence of the two parameters involved in the procedure to avoid the prediction of small ligand binding pockets on PPV values.** The $CV_{local}{}^{high}$ (x-axis) and the percentage of residues allowed to show $CV_{local} \geq CV_{local}{}^{high}$ (y-axis) are the two parameters involved in the procedure to avoid the prediction of small ligand binding pockets. In each cell is reported the averaged PPV value computed on the HR-PDNA187 dataset. Two different sets of $score_{Clus}^{Seed}$, $score_{Clus}^{ExtOut}$, $score_{Res}^{Layer=seed,extension,outer\ layer}$ and $Confidence_{ClusterSize}$ were tried, reported on the left and right columns respectively, to test the stability of results with respect to these other thresholds used in the algorithm. In each column, three 2D plots are reported for predictions obtained with a consensus of 2, 5 and 8 runs out of 10, respectively. Default values chosen in $JET_{DNA}^2$ for D-SC1 and D-SC2 are highlighted in yellow and orange, respectively.

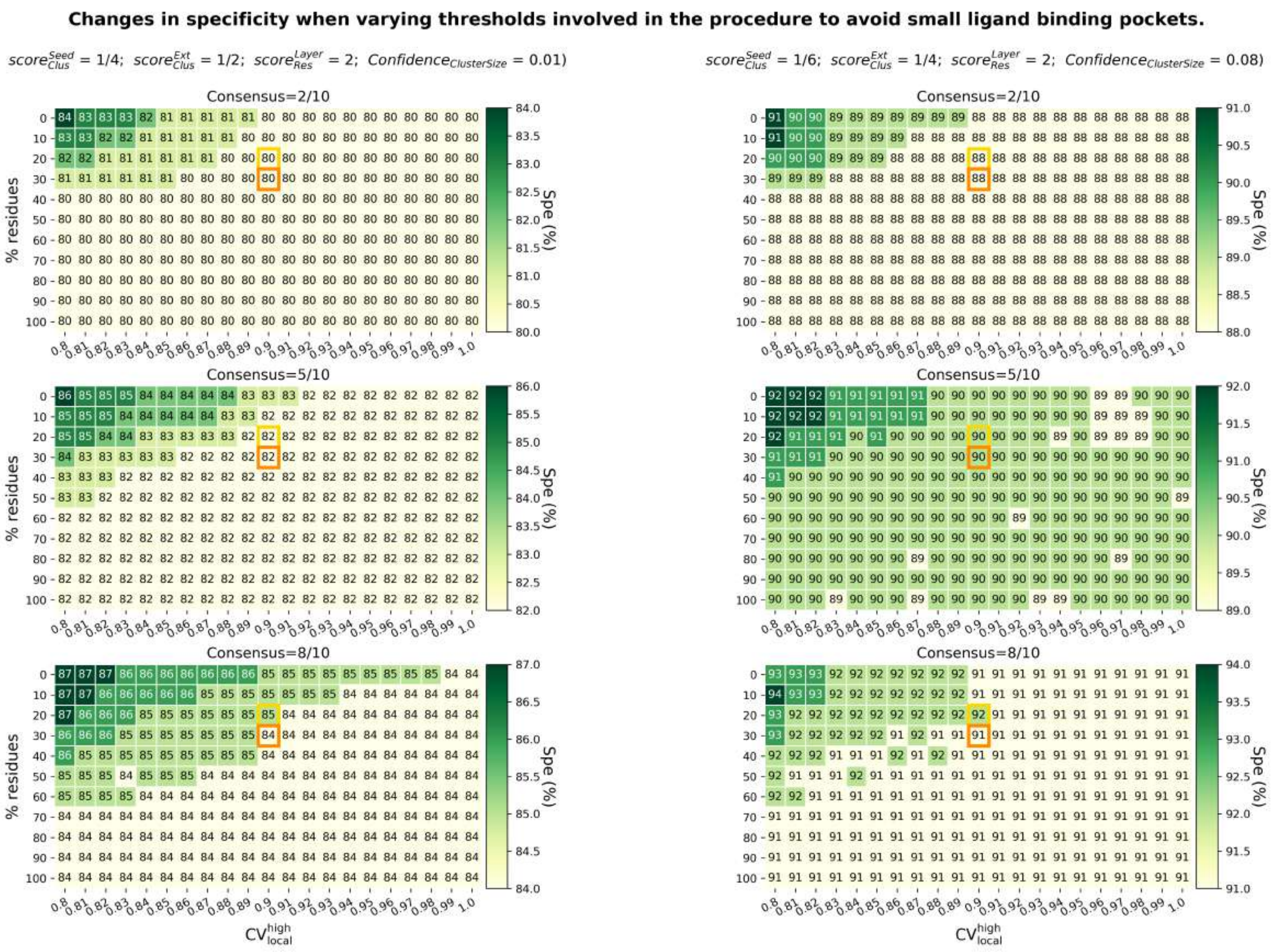


Figure S14: **Influence of the two parameters involved in the procedure to avoid the prediction of small ligand binding pockets on specificity values.** The $CV_{local}{}^{high}$ (x-axis) and the percentage of residues allowed to show $CV_{local} \geq CV_{local}{}^{high}$ (y-axis) are the two parameters involved in the procedure to avoid the prediction of small ligand binding pockets. In each cell is reported the averaged specificity value computed on the HR-PDNA187 dataset. Two different sets of $score_{Clus}^{Seed}$, $score_{Clus}^{ExtOut}$, $score_{Res}^{Layer=seed,extension,outer\ layer}$ and $Confidence_{ClusterSize}$ were tried, reported on the left and right columns respectively, to test the stability of results with respect to these other thresholds used in the algorithm. In each column, three 2D plots are reported for predictions obtained with a consensus of 2, 5 and 8 runs out of 10, respectively. Default values chosen in $JET_{DNA}^2$ for D-SC1 and D-SC2 are highlighted in yellow and orange, respectively.

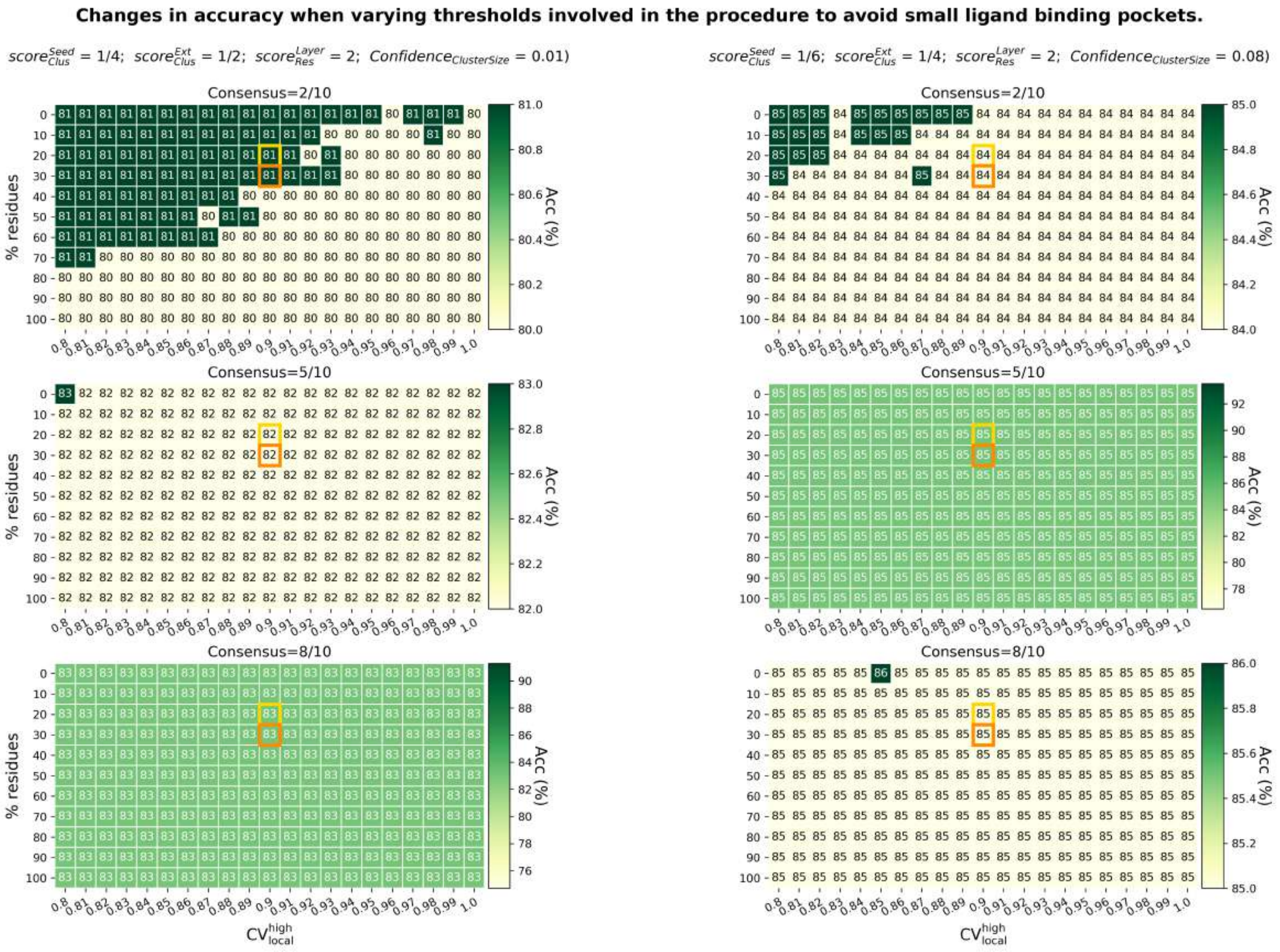


Figure S15: **Influence of the two parameters involved in the procedure to avoid the prediction of small ligand binding pockets on accuracy values.** The $CV_{local}{}^{high}$ (x-axis) and the percentage of residues allowed to show $CV_{local} \geq CV_{local}{}^{high}$ (y-axis) are the two parameters involved in the procedure to avoid the prediction of small ligand binding pockets. In each cell is reported the averaged accuracy value computed on the HR-PDNA187 dataset. Two different sets of $score_{Clus}^{Seed}$, $score_{Clus}^{ExtOut}$, $score_{Res}^{Layer=seed,extension,outer\ layer}$ and $Confidence_{ClusterSize}$ were tried, reported on the left and right columns respectively, to test the stability of results with respect to these other thresholds used in the algorithm. In each column, three 2D plots are reported for predictions obtained with a consensus of 2, 5 and 8 runs out of 10, respectively. Default values chosen in $JET^2_{DNA}$ for D-SC1 and D-SC2 are highlighted in yellow and orange, respectively.

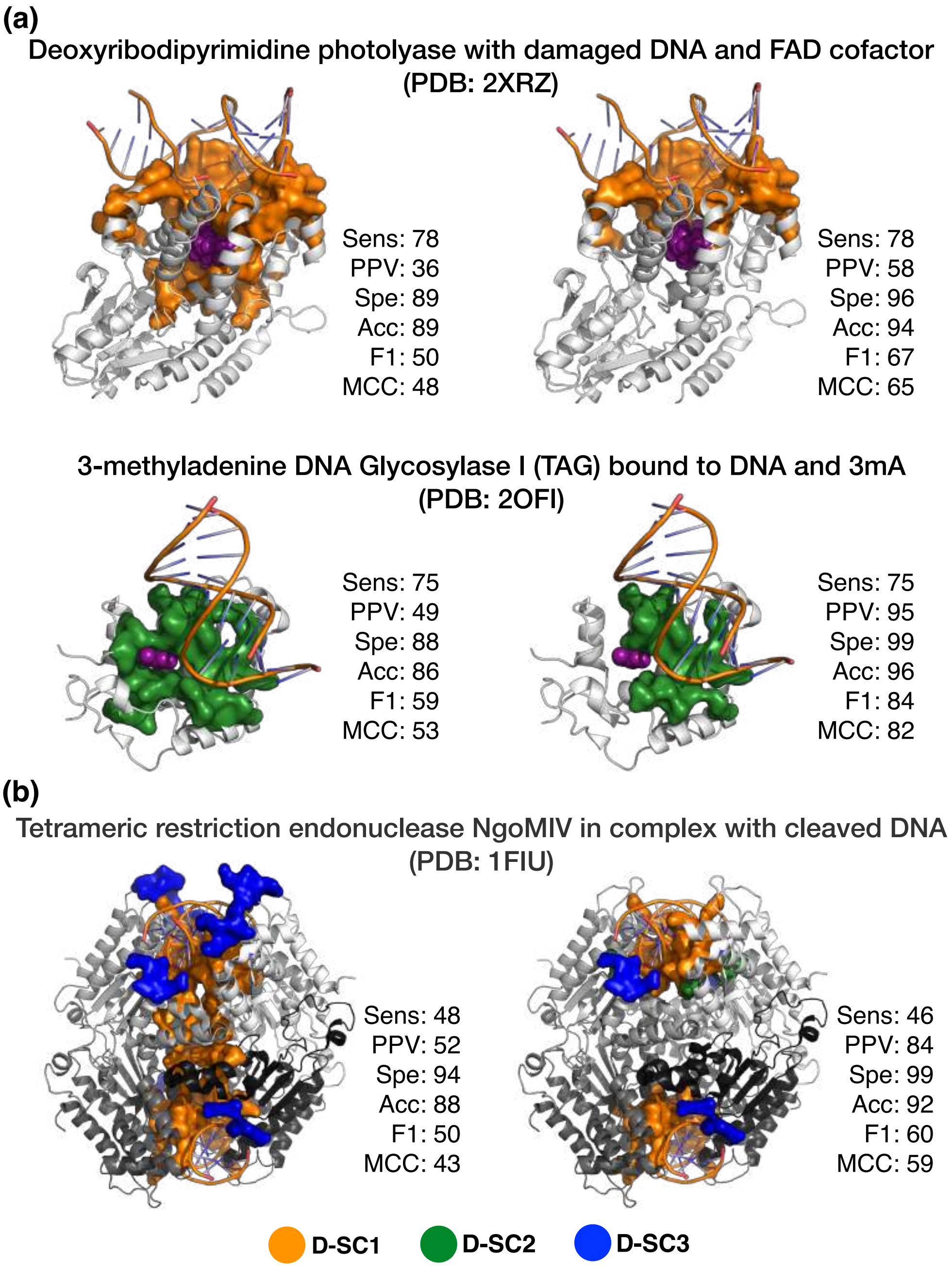


Figure S16: **Examples of improvements in $\mathrm{JET}^2_{\mathrm{DNA}}$ predictions by the implementation of the procedure to avoid small ligand binding pockets.** On the left, $\mathrm{JET}^2_{\mathrm{DNA}}$ predictions without the procedure to avoid small ligand binding pockets. On the right, $\mathrm{JET}^2_{\mathrm{DNA}}$ predictions with the procedure using the default values reported in the paper ($\mathrm{CV}_{\mathrm{local}} = 0.9$ and $\%\ residues_{allowed} = 20\%$ (D-SC1), 30% (D-SC2)). (a) Two examples showing an improvement in the $\mathrm{JET}^2_{\mathrm{DNA}}$ prediction for both D-SC1 and D-SC2 by avoiding the prediction of the small ligand binding pocket close to the DNA-binding site. On top, a deoxyribodipyrimidine photolyase in complex with a damaged duplex DNA and a FAD cofactor (represented with purple spheres) (PDB code: 2XRZ). On bottom, the 3-methyladenine DNA Glycosylase I (TAG) in complex with abasic duplex DNA and a 3mA nucleobase (represented with purple spheres) (PDB code: 2OFI). (b) An example showing the procedure implemented to avoid small ligand binding pockets can ameliorate predictions on multimeric protein structures. The tetrameric restriction endonuclease NgoMIV in complex with two duplex DNA (PDB code: 1FIU) is reported with different protein chains greyscale colored. $\mathrm{JET}^2_{\mathrm{DNA}}$ predictions by D-SC1, D-SC2 and D-SC3 are represented as orange, green and blue opaque colored surfaces, respectively. All the predictions were obtained from a consensus of 2 runs out of 10 of $\mathrm{JET}^2_{\mathrm{DNA}}$.

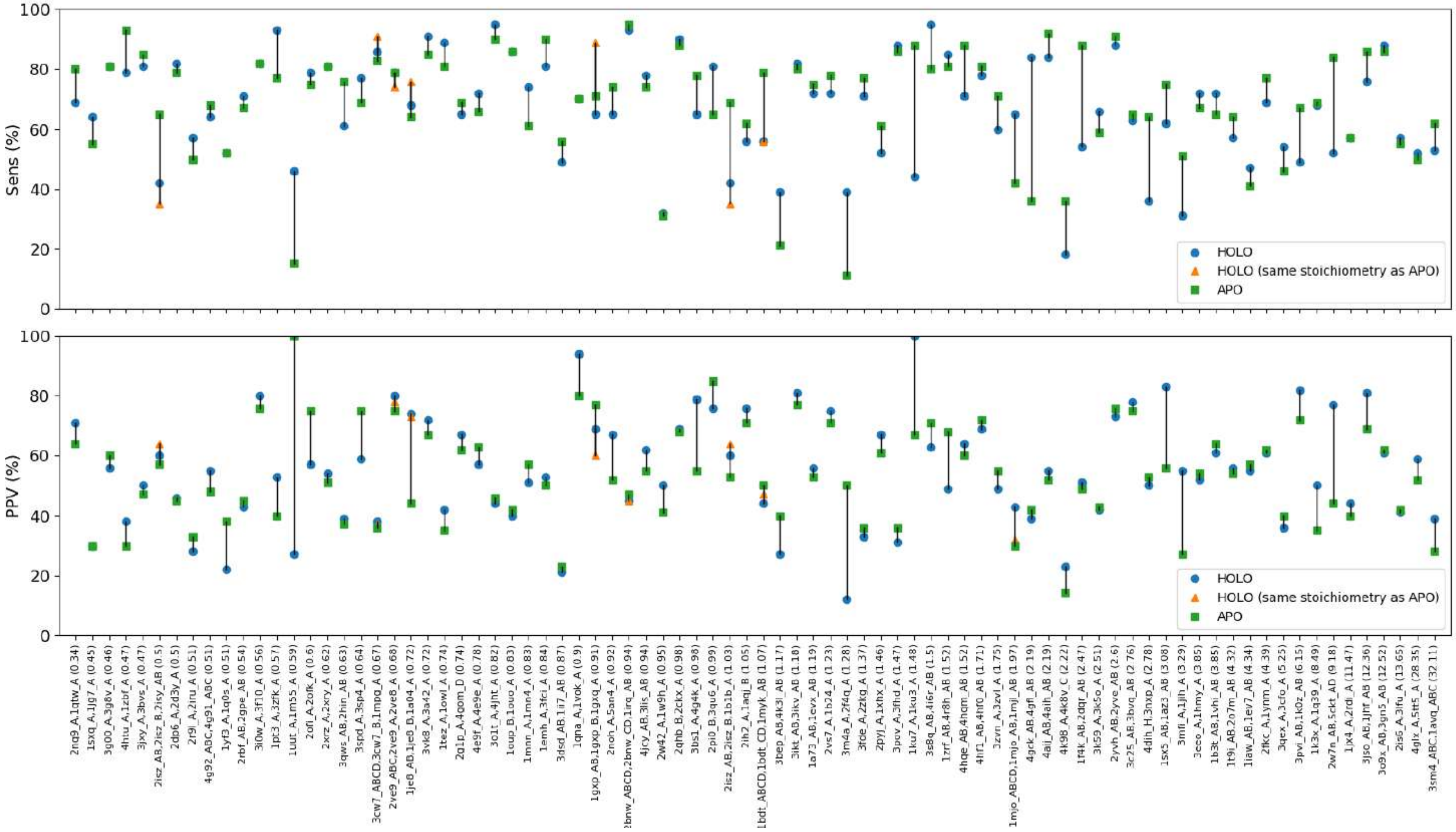


Figure S17: **Variation in sensitivity and PPV values between predictions on HOLO and APO forms.** On the x-axis are reported the 82 HOLO-APO pairs (HOLO-APO82 dataset), ordered by the increasing RMSD between bound and unbound conformations of each pair. For the 8 pairs showing a HOLO form in a different stoichiometry (more protein chains) than the APO one, the subset of chains in the HOLO form corresponding to the same stoichiometry of the APO one is also reported. The format for the 74 HOLO-APO pairs in the same stoichiometry is "HOLO, APO (RMSD HOLO-APO)", while the format for the 8 HOLO-APO pairs in a different stoichiometry is "HOLO, HOLO_subsetChains, APO (RMSD HOLO-APO)". On the y-axis, sensitivity and PPV values are reported in percentage. Performance on HOLO, HOLO_subsetChains and APO forms are reported in blue circles, orange triangles and green squares, respectively. $iJET^2_{DNA}$ predictions were obtained with a consensus of 2 runs out of 10.

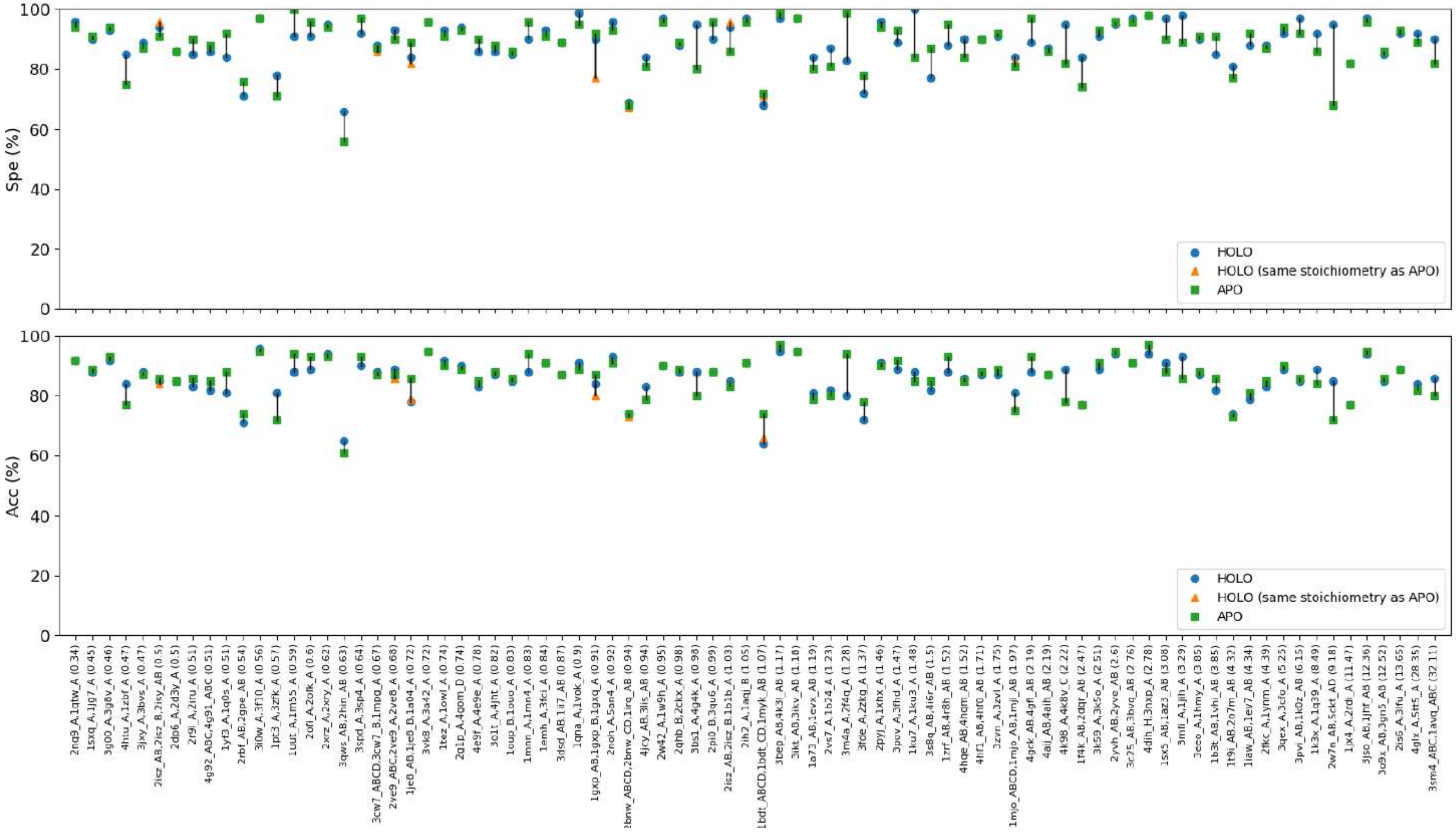


Figure S18: **Variation in specificity and accuracy between predictions on HOLO and APO forms.** On the x-axis are reported the 82 HOLO-APO pairs (HOLO-APO82 dataset), ordered by the increasing RMSD between bound and unbound conformations of each pair. For the 8 pairs showing a HOLO form in a different stoichiometry (more protein chains) than the APO one, the subset of chains in the HOLO form corresponding to the same stoichiometry of the APO one is also reported. The format for the 74 HOLO-APO pairs in the same stoichiometry is "HOLO, APO (RMSD HOLO-APO)", while the format for the 8 HOLO-APO pairs in a different stoichiometry is "HOLO, HOLO_subsetChains, APO (RMSD HOLO-APO)". On the y-axis, specificity and accuracy values are reported in percentage. Performance on HOLO, HOLO_subsetChains and APO forms are reported in blue circles, orange triangles and green squares, respectively. $\text{iJET}^2_{\text{DNA}}$ predictions were obtained with a consensus of 2 runs out of 10.

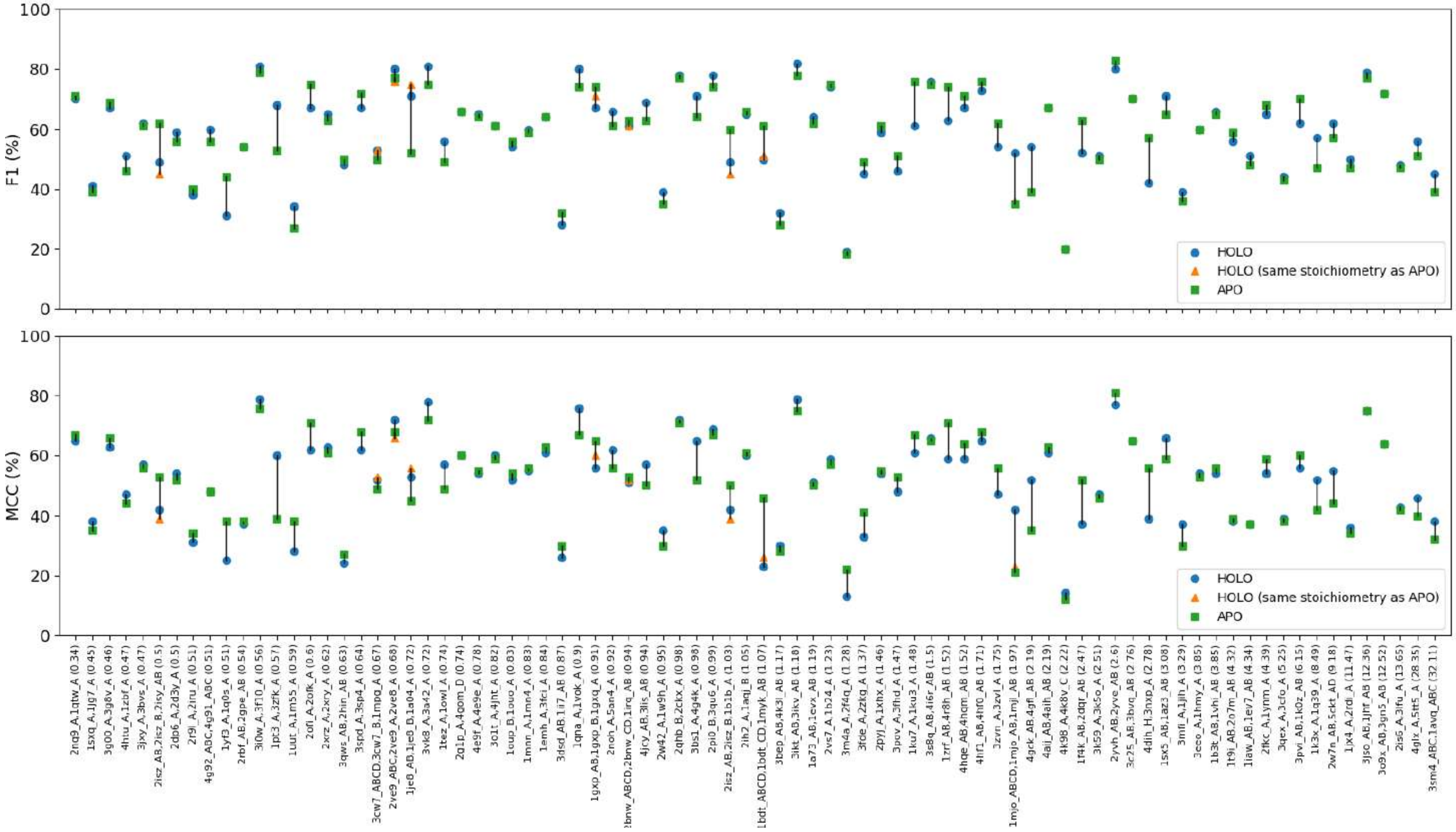


Figure S19: **Variation in F1 and MCC values between predictions on HOLO and APO forms.** On the x-axis are reported the 82 HOLO-APO pairs (HOLO-APO82 dataset), ordered by the increasing RMSD between bound and unbound conformations of each pair. For the 8 pairs showing a HOLO form in a different stoichiometry (more protein chains) than the APO one, the subset of chains in the HOLO form corresponding to the same stoichiometry of the APO one is also reported. The format for the 74 HOLO-APO pairs in the same stoichiometry is "HOLO, APO (RMSD HOLO-APO)", while the format for the 8 HOLO-APO pairs in a different stoichiometry is "HOLO, HOLO_subsetChains, APO (RMSD HOLO-APO)". On the y-axis, F1 and MCC values are reported in percentage. Performance on HOLO, HOLO_subsetChains and APO forms are reported in blue circles, orange triangles and green squares, respectively. iJET$^2_{DNA}$ predictions were obtained with a consensus of 2 runs out of 10.

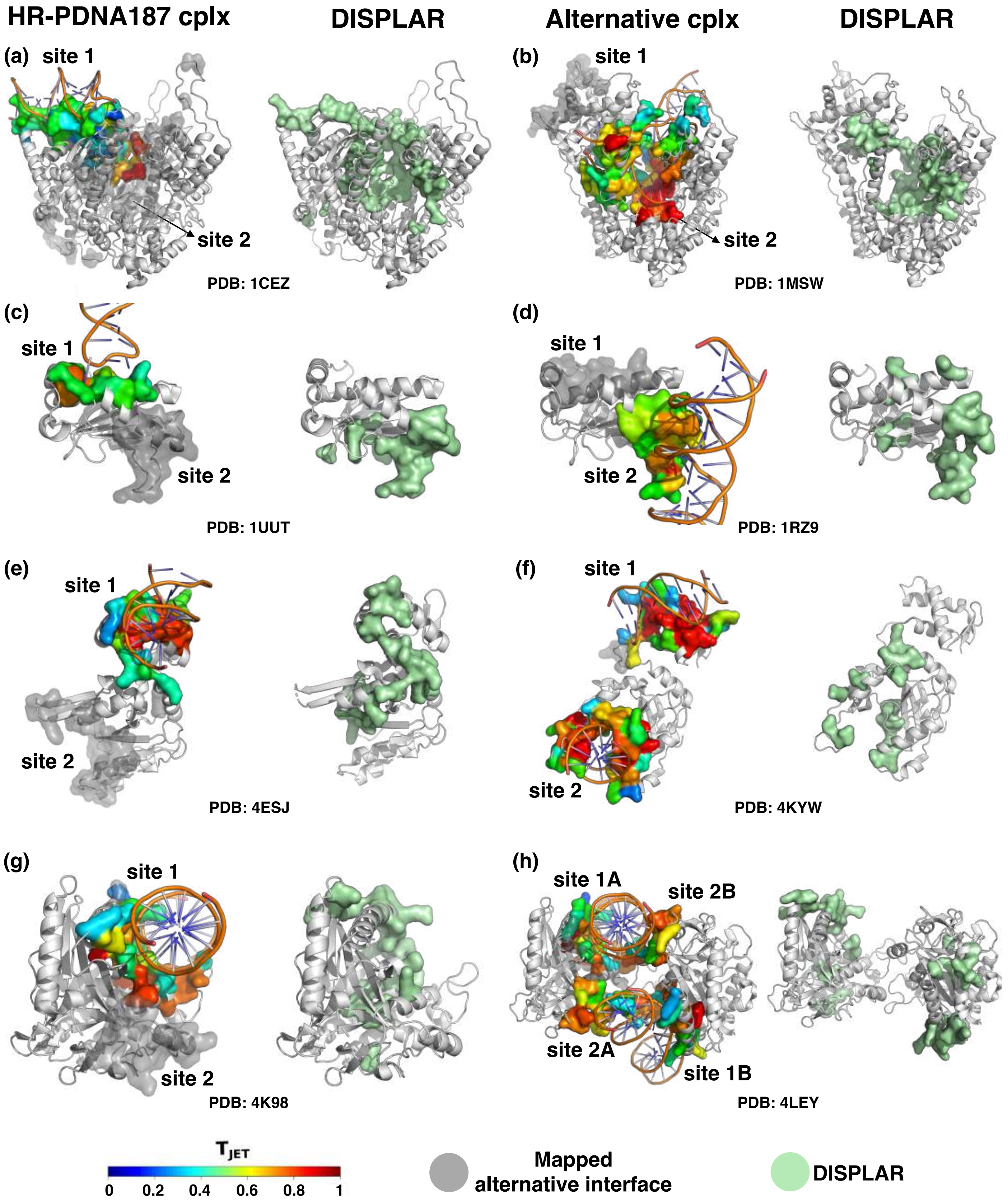


Figure S20: **DISPLAR predictions on proteins presenting multiple DNA-binding sites.** DISPLAR predicted patches were defined as formed by residue indicated as predicted in the tool results and are colored in lightgreen. Compare with Fig. 5.

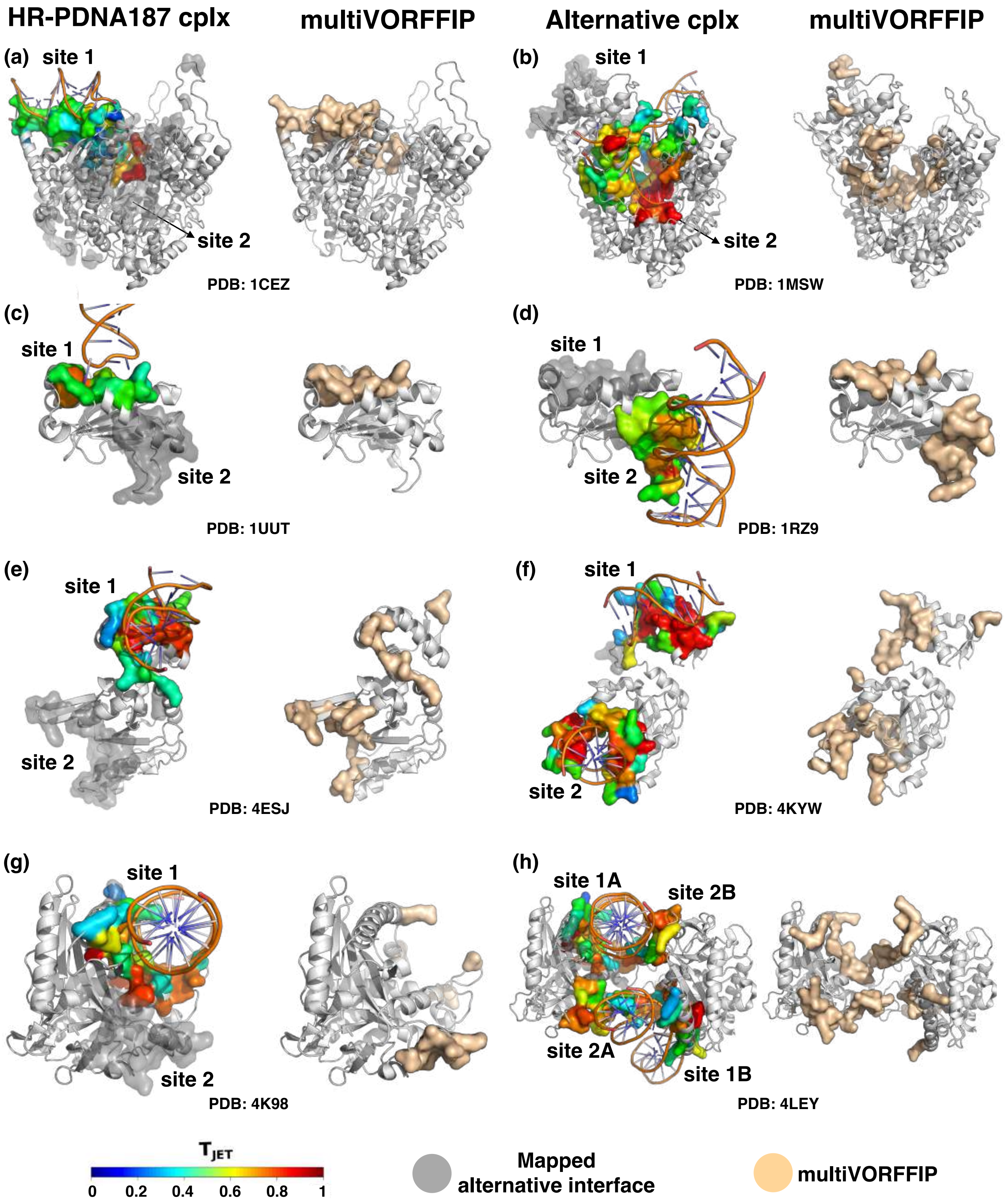


Figure S21: **multiVORFFIP predictions on proteins presenting multiple DNA-binding sites.** multiVORFFIP predicted patches were defined as formed by residues with probability > 0.5 and are colored in beige. Compare with Fig. 5.

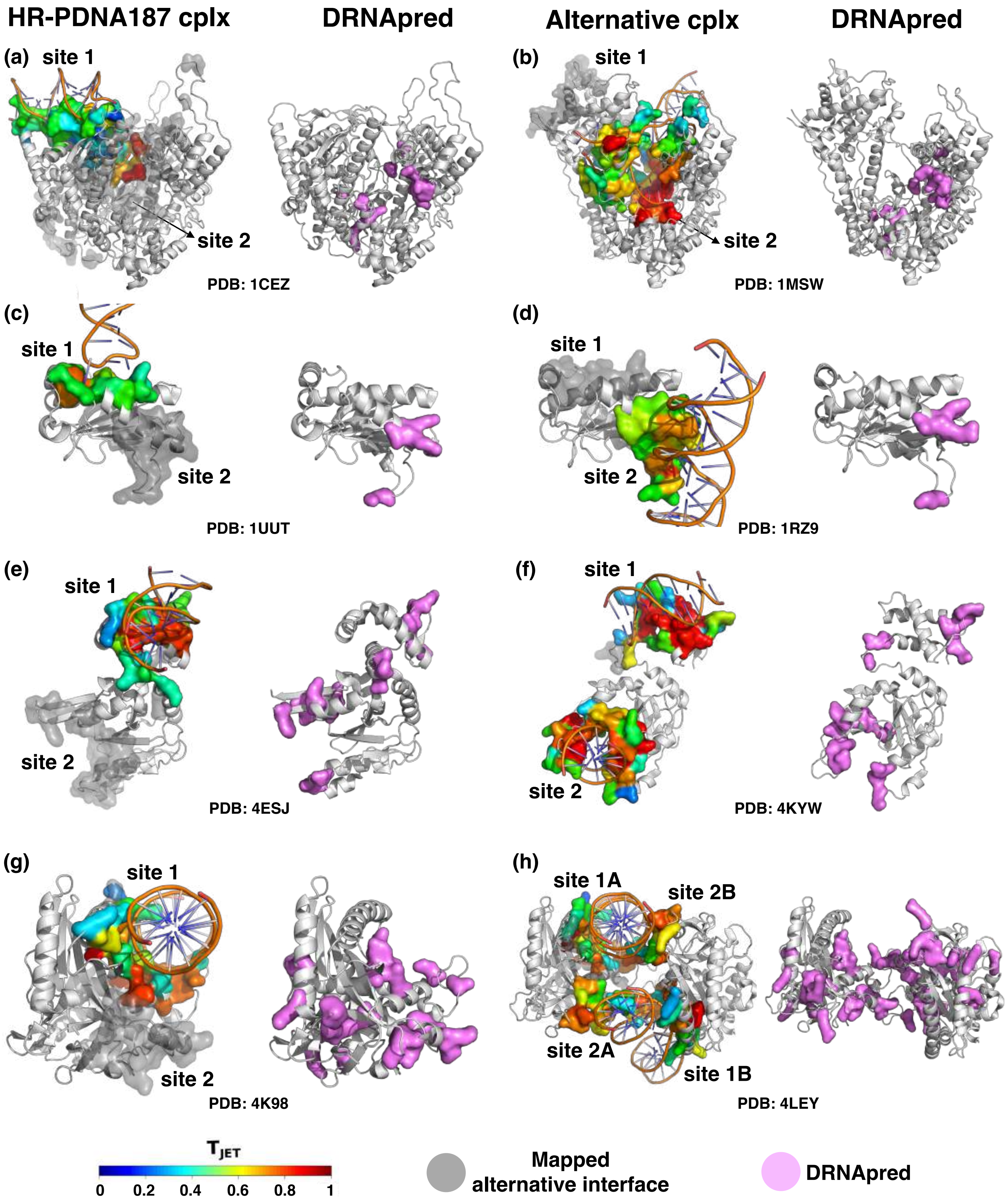


Figure S22: **DRNApred predictions on proteins presenting multiple DNA-binding sites.** DRNApred predicted patches were defined as formed by residues indicated as predicted by the binary prediction column in the tool results and are colored in orchid. Compare with Fig. 5.
