## Supplementary Tables for "Multiple protein-DNA interfaces unravelled by evolutionary information, physico-chemical and geometrical properties"

Table S1: **List of the 187 complexes comprised in HR-PDNA187 dataset and the 82 HOLO-APO pairs available.** The entries in the columns are respectively: 1) the Protein Data Bank (PDB) [1] identifier of the HR-PDNA187 complex; 2) the protein chains considered in the complex; 3) the subset of protein chains reported in column 2 non redundant at 95% of sequence identity; 4) the DNA chains considered in the complex; 5) the PDB identifier and the considered chains of the HOLO form in the HOLO-APO82 dataset; 6) the PDB identifier and the considered chains of the APO form in the HOLO-APO82 dataset; 7) the class and 8) the subclass to which the protein belongs as derived from the Nucleic Acid Database [2]. Concerning the protein stoichiometry, the HR-PDNA187 comprises: 109 monomers, 64 homo-2-mers, 3 hetero-2-mers, 3 homo-3-mers, 1 hetero-3-mer, 5 homo-4-mers, 1 hetero-5-mer and 1 homo-6-mer. The APO forms in the HOLO-APO82 dataset are divided in: 52 monomers, 28 homo-2-mers, 1 homo-3-mer and 1 hetero-3-mer. Specifically, we observed a change in the stoichiometry in 4 HOLO-APO pairs: 3 homo-4-mers, 1 homo-4-mer, 1 homo-3-mer, 1 homo-2-mer in the bound form are respectively 3 homo-2-mers and 3 monomers in the unbound form.

| PDB ID | prot chains | prot nr chains | DNA chains | HOLO ID:c | APO ID:c | Class | Subclass |
|---|---|---|---|---|---|---|---|
| 1a3q | AB | A | CD | | | regulatory | transcription factor |
| 1a73 | AB | A | CDEF | 1a73:BA | 1evx:AB | enzyme | nuclease |
| 1b3t | AB | A | CD | 1b3t:BA | 1vhi:AB | regulatory | other |
| 1bdt | ABCD | A | EF | 1bdt:CD | 1myk:AB | regulatory | gene |
| 1bl0 | A | A | BC | | | regulatory | transcription factor |
| 1cez | A | A | NT | | | enzyme | polymerase |
| 1d02 | AB | A | CD | | | enzyme | nuclease |
| 1dc1 | AB | A | CW | | | enzyme | nuclease |
| 1dfm | AB | A | CD | | | enzyme | nuclease |
| 1egw | AB | A | EF | | | regulatory | transcription factor |
| 1emh | A | A | BC | 1emh:A | 3fci:A | enzyme | glycosylase |
| 1esg | AB | A | CD | | | enzyme | nuclease |
| 1f4k | AB | A | DE | 1f4k:BA | 2dqr:AB | regulatory | replication |
| 1fiu | ABCD | A | EFGHIJKL | | | enzyme | nuclease |
| 1gu4 | AB | A | CD | | | regulatory | transcription factor |
| 1gxp | AB | A | CD | 1gxp:B | 1gxq:A | regulatory | other |
| 1h6f | AB | A | CD | | | regulatory | transcription factor |
| 1hlv | A | A | BC | | | structural | other/centromere |
| 1i3j | A | A | BC | | | enzyme | nuclease |
| 1iaw | AB | A | CDEF | 1iaw:BA | 1ev7:AB | enzyme | hydrolase |
| 1j3e | A | A | BC | | | regulatory | replication |
| 1je8 | AB | A | CD | 1je8:B | 1a04:A | regulatory | transcription factor |
| 1jko | C | C | AB | | | enzyme | recombinase |
| 1jx4 | A | A | PT | 1jx4:A | 2rdi:A | enzyme | polymerase |
| 1k3x | A | A | BC | 1k3x:A | 1q39:A | enzyme | nuclease |
| 1k4t | A | A | BCD | | | enzyme | isomerase |
| 1ku7 | A | A | BC | 1ku7:A | 1ku3:A | regulatory | transcription factor |
| 1kx5 | ABCDEFGH | B<br>D<br>A<br>C | IJ | | | structural | histone |
| 1l3l | BD | B | EG | | | regulatory | transcription factor |
| 1lmb | 34 | 3 | 12 | | | regulatory | other |
| 1lq1 | CD | C | EF | | | regulatory | transcription factor |
| 1mjo | ABCD | A | FG | 1mjo:AB | 1mjl:AB | regulatory | transcription factor |
| 1mnn | A | A | BC | 1mnn:A | 1mn4:A | regulatory | transcription factor |
| 1nkp | AB | B<br>A | FG | | | regulatory | transcription factor |
| 1oe4 | AB | A | EF | | | enzyme | glycosylase |
| 1orn | A | A | BC | | | enzyme | nuclease |
| 1oup | B | B | CD | 1oup:B | 1ouo:A | enzyme | nuclease |
| 1owf | AB | A<br>B | CDE | | | regulatory | transcription factor |
| 1ozj | A | A | CD | | | regulatory | transcription factor |

Continued on next page

Table S1 – continued from previous page

| PDB ID | prot chains | prot nr chains | DNA chains | HOLO ID:c | APO ID:c | Class | Subclass |
|---|---|---|---|---|---|---|---|
| 1pp7 | U | U | EF | | | regulatory | transcription factor |
| 1pt3 | A | A | CDEFGH | 1pt3:A | 3zfk:A | enzyme | nuclease |
| 1qna | A | A | CD | 1qna:A | 1vok:A | regulatory | transcription factor |
| 1r71 | AB | A | EFIJ | | | regulatory | transcription factor |
| 1rh6 | B | B | CD | | | regulatory | recombination |
| 1rxw | A | A | BC | | | enzyme | nuclease |
| 1sa3 | A | A | CD | | | enzyme | nuclease |
| 1skn | P | P | AB | | | regulatory | transcription factor |
| 1sx5 | AB | A | CDEF | 1sx5:AB | 1az3:AB | enzyme | nuclease |
| 1sxq | A | A | CE | 1sxq:A | 1jg7:A | enzyme | transferase |
| 1t7p | A | A | PT | | | enzyme | polymerase |
| 1t9i | AB | A | CD | 1t9i:AB | 2o7m:AB | enzyme | nuclease |
| 1tc3 | C | C | AB | | | enzyme | other |
| 1tez | A | A | IJK | 1tez:A | 1owl:A | enzyme | lyase |
| 1u8b | A | A | BCDE | | | regulatory | other |
| 1uut | A | A | C | 1uut:A | 1m55:A | enzyme | nuclease |
| 1wb9 | AB | A | EF | | | regulatory | repair |
| 1xyi | A | A | BC | | | structural | chromosomal |
| 1yf3 | A | A | CD | 1yf3:A | 1q0s:A | enzyme | methyl |
| 1yo5 | C | C | AB | | | regulatory | transcription factor |
| 1zme | CD | C | AB | | | regulatory | transcription factor |
| 1zrf | AB | A | WXYZ | 1zrf:AB | 4r8h:AB | regulatory | other |
| 2aor | A | A | CD | | | enzyme | methyl |
| 2aq4 | A | A | PT | | | enzyme | transferase |
| 2bnw | ABCD | A | EFGH | 2bnw:CD | 1irq:AB | regulatory | other |
| 2dp6 | A | A | CD | 2dp6:A | 2d3y:A | enzyme | glycosylase |
| 2e52 | AB | A | EG | | | enzyme | nuclease |
| 2ex5 | AB | A | XY | | | enzyme | nuclease |
| 2fkc | A | A | CD | 2fkc:A | 1ynm:A | enzyme | nuclease |
| 2g1p | A | A | FG | 2g1p:A | 4gom:D | enzyme | methyl |
| 2gb7 | AB | A | EF | | | enzyme | nuclease |
| 2h27 | A | A | BC | | | enzyme | transferase |
| 2h7g | X | X | YZ | | | enzyme | isomerase |
| 2i06 | A | A | BC | | | regulatory | replication |
| 2ih2 | A | A | BC | 2ih2:A | 1aqj:B | enzyme | methyl |
| 2ihm | A | A | DPT | | | enzyme | polymerase |
| 2is6 | A | A | CD | 2is6:A | 3lfu:A | enzyme | helicase |
| 2isz | AB | A | EF | 2isz:BA<br>2isz:B | 2isy:AB<br>1b1b:A | regulatory | transcription factor |
| 2noh | A | A | BC | 2noh:A | 5an4:A | enzyme | glycosylase |
| 2nq9 | A | A | BCD | 2nq9:A | 1qtw:A | enzyme | nuclease |
| 2o4a | A | A | BC | | | regulatory | transcription factor |
| 2ofi | A | A | BC | 2ofi:A | 2ofk:A | enzyme | glycosylase |
| 2pi0 | B | B | EF | 2pi0:B | 3qu6:A | regulatory | other |
| 2pyj | A | A | XY | 2pyj:A | 1xhx:A | enzyme | polymerase |
| 2qhb | B | B | CD | 2qhb:B | 2ckx:A | structural | telomere |
| 2qoj | Z | Z | XY | | | enzyme | nuclease |
| 2r1j | LR | L | AB | | | regulatory | transcription factor |
| 2r9l | A | A | CD | 2r9l:A | 2iru:A | enzyme | polymerase |
| 2rbf | AB | A | CD | 2rbf:BA | 2gpe:AB | regulatory | other |
| 2ve9 | ABC | A | IJ | 2ve9:A | 2ve8:A | structural | other |
| 2vla | A | A | LM | | | enzyme | nuclease |
| 2vs7 | A | A | BC | 2vs7:A | 1b24:A | enzyme | nuclease |
| 2w42 | A | A | PQ | 2w42:A | 1w9h:A | regulatory | other |
| 2w7n | AB | A | EFGH | 2w7n:BA | 5ckt:AD | regulatory | gene |
| 2xm3 | CD | C | KLMN | | | enzyme | other/transposase |
| 2xrz | A | A | CD | 2xrz:A | 2xry:A | enzyme | lyase |
| 2xzf | A | A | BC | | | enzyme | glycosylase |

Continued on next page

**Table S1 – continued from previous page**

| PDB ID | prot chains | prot nr chains | DNA chains | HOLO ID:c | APO ID:c | Class | Subclass |
|---|---|---|---|---|---|---|---|
| 2yvh | AB | A | EFGH | 2yvh:AB | 2yve:AB | regulatory | transcription factor |
| 3aaf | A | A | CD | | | enzyme | other |
| 3bep | AB | A | CD | 3bep:BA | 4k3l:AB | enzyme | polymerase |
| 3bm3 | AB | A | CD | | | enzyme | nuclease |
| 3bs1 | A | A | BC | 3bs1:A | 4g4k:A | regulatory | gene |
| 3c0w | A | A | BCD | | | enzyme | nuclease |
| 3c25 | AB | A | CD | 3c25:AB | 3bvq:AB | enzyme | nuclease |
| 3coq | AB | A | DE | | | regulatory | transcription factor |
| 3cw7 | ABCD | A | EFGH | 3cw7:B | 1mpg:A | enzyme | glycosylase |
| 3dsd | AB | A | C | 3dsd:BA | 1ii7:AB | regulatory | repair |
| 3dvo | AB | A | EF | | | enzyme | nuclease |
| 3eeo | A | A | CD | 3eeo:A | 1hmy:A | enzyme | methyl |
| 3f2b | A | A | PT | | | enzyme | polymerase |
| 3fde | A | A | DE | 3fde:A | 2zkg:A | enzyme | ligase |
| 3fdq | AB | A | CD | | | regulatory | other |
| 3g00 | A | A | HI | 3g00:A | 3g8v:A | enzyme | nuclease |
| 3g0q | A | A | BC | | | enzyme | hydrolase |
| 3g9m | AB | A | CD | | | regulatory | transcription factor |
| 3gox | AB | A | CD | | | enzyme | nuclease |
| 3gxq | AB | A | CD | | | regulatory | other |
| 3h0d | AB | A | CD | | | regulatory | transcription factor |
| 3i0w | A | A | BC | 3i0w:A | 3f10:A | enzyme | glycosylase |
| 3iag | C | C | AB | | | regulatory | transcription factor |
| 3iay | A | A | PT | | | enzyme | polymerase |
| 3igm | AB | A | CDWX | | | regulatory | transcription factor |
| 3ikt | AB | A | CD | 3ikt:AB | 3ikv:AB | regulatory | other |
| 3jso | AB | A | CD | 3jso:AB | 1jhf:AB | regulatory | other |
| 3jxy | A | A | BC | 3jxy:A | 3bvs:A | enzyme | glycosylase |
| 3k59 | A | A | PT | 3k59:A | 3k5o:A | enzyme | polymerase |
| 3kde | C | C | AB | | | enzyme | other |
| 3kxt | A | A | BC | | | structural | other |
| 3l2c | A | A | BC | | | regulatory | transcription factor |
| 3lap | ABCDEF | A | GHIJKL | | | regulatory | other |
| 3m4a | A | A | DE | 3m4a:A | 2f4q:A | enzyme | isomerase |
| 3mfi | A | A | PT | 3mfi:A | 1jih:A | enzyme | polymerase |
| 3mln | AB | A | CD | | | regulatory | transcription factor |
| 3mva | O | O | DE | | | regulatory | transcription factor |
| 3mx4 | AH | A | KL | | | enzyme | nuclease |
| 3o1t | A | A | BC | 3o1t:A | 4jht:A | enzyme | other |
| 3o9x | AB | A | EF | 3o9x:AB | 3gn5:AB | regulatory | gene |
| 3od8 | A | A | IJ | | | enzyme | other |
| 3pov | A | A | CD | 3pov:A | 3fhd:A | enzyme | other |
| 3pvi | AB | A | CD | 3pvi:AB | 1k0z:AB | enzyme | nuclease |
| 3pvv | A | A | CD | | | regulatory | replication |
| 3qex | A | A | PT | 3qex:A | 3cfo:A | enzyme | polymerase |
| 3qmd | A | A | BC | | | regulatory | other |
| 3qqy | A | A | BC | | | enzyme | nuclease |
| 3qws | AB | A | CN | 3qws:AB | 2hin:AB | regulatory | other |
| 3rkq | A | A | CD | | | regulatory | transcription factor |
| 3rmp | AC | A | EFGH | | | enzyme | other |
| 3s57 | A | A | BC | | | enzyme | other |
| 3s8q | AB | A | CD | 3s8q:BA | 4i6r:AB | regulatory | other |
| 3sjm | A | A | CD | | | structural | telomere |
| 3sm4 | ABC | A | DE | 3sm4:CAB | 1avq:ABC | enzyme | nuclease |
| 3spd | A | A | EF | 3spd:A | 3sp4:A | enzyme | hydrolase |
| 3ssc | A | A | CD | | | enzyme | nuclease |
| 3tan | A | A | BC | | | enzyme | polymerase |
| 3tq6 | A | A | CD | | | regulatory | transcription factor |

Continued on next page

Table S1 – continued from previous page

| PDB ID | prot chains | prot nr chains | DNA chains | HOLO ID:c | APO ID:c | Class | Subclass |
|---|---|---|---|---|---|---|---|
| 3u2b | C | C | AB | | | regulatory | transcription factor |
| 3vk8 | A | A | CD | 3vk8:A | 3a42:A | enzyme | glycosylase |
| 3vxv | A | A | BC | | | enzyme | hydrolase |
| 3zvk | FG | E | XY | | | regulatory | other |
| 3zvn | A | A | EFGHI | 3zvn:A | 3zvl:A | enzyme | hydrolase |
| 4aij | AB | A | CD | 4aij:BA | 4aih:AB | regulatory | transcription factor |
| 4dih | H | H | D | 4dih:H | 3nxp:A | enzyme | thrombin |
| 4e9f | A | A | CD | 4e9f:A | 4e9e:A | enzyme | Hydrolase/glycosylase |
| 4ecq | A | A | PT | | | enzyme | polymerase |
| 4esj | A | A | CD | | | enzyme | nuclease |
| 4fzx | C | C | AB | | | enzyme | nuclease |
| 4g92 | ABC | B C A | DE | 4g92:ABC | 4g91:ABC | regulatory | transcription factor |
| 4gck | AB | A | WZ | 4gck:AB | 4gfl:AB | other | other |
| 4gjr | AB | A | GHIJ | | | regulatory | transcription factor |
| 4glx | A | A | BCD | 4glx:A | 5tt5:A | enzyme | ligase |
| 4gzn | C | C | AB | | | regulatory | transcription factor |
| 4h0e | B | B | TU | | | regulatory | transcription factor |
| 4h10 | AB | B A | CD | | | regulatory | transcription factor |
| 4hf1 | AB | A | CD | 4hf1:AB | 4hf0:AB | regulatory | transcription factor |
| 4hqe | AB | A | CD | 4hqe:BA | 4hqm:AB | regulatory | transcription factor |
| 4htu | A | A | CD | 4htu:A | 1zbf:A | enzyme | nuclease |
| 4i2o | AB | A | XW | | | regulatory | other |
| 4ix7 | AB | A | CD | | | regulatory | other |
| 4j3n | AB | A | CDEF | | | enzyme | isomerase |
| 4jbm | A | A | RT | | | regulatory | other |
| 4jcy | AB | A | CD | 4jcy:BA | 3lis:AB | regulatory | other |
| 4k98 | A | A | DE | 4k98:A | 4k8v:C | enzyme | transferase |
| 4kb1 | A | A | C | | | enzyme | hydrolase |
| 4kli | A | A | DPT | | | enzyme | polymerase |
| 4kpy | A | A | CDN | | | NoClass | NoClass |
| 4qtj | A | A | BC | | | regulatory | transcription factor |
| 4rkh | CEF | C | AB | | | enzyme | ligase |
| 6pax | A | A | BC | | | regulatory | transcription factor |

Table S2: **$\mathbf{JET^2_{DNA}}$ algorithm**

$R \leftarrow \{r_i, score(r_i) > score^{seed}_{res}\}$;
**for** $r_i \in R$ *and* $\notin C$ **do**
**if** $score(r_i) > score^{seed}_{clus}$ **then**
$newClus \leftarrow r_i$;
**while** $\{neighbors\ of\ newClus\}_{\in R} \neq \{\}$ **do**
**for** $r_j \in \{neighbors\ of\ newClus\}_{\in R}$ **do**
**if** $\mu(newClus + r_j) > score^{seed}_{clus}$ **then**
add $r_j$ to $newClus$;
**end**
**end**
**end**
**end**
**end**
**if** $r_i \in C, CV_{local}(r_i) > 0.9 / r_i \in C > threshold_{buried}$ **then**
remove all residues $r_i, CV_{local}(r_i) > 0.9$ from $R$ and restart clustering;
**end**
merge clusters < 5Å away from each other;

**Seeds detection**

$C$, ensemble of detected clusters

$score(r_i)$, score of the residue $r_i$

$\mu(newClus + r_j)$, mean score computed over the ensemble of residues of cluster $newClus$ and $r_j$

$score^{seed}_{res}$, threshold score for residues during seeds detection

$score^{seed}_{clus}$, threshold score for clusters during seeds detection

$\{neighbors\ of\ newClus\}$, ensemble of residues < 5Å away from $newClus$

$r_i \in C, CV_{local}(r_i) > 0.9 | / | r_i \in C$, percentage of residues in detected clusters with $CV_{local} > 0.9$

$threshold_{buried}$, maximum percentage of residues with $CV_{local} > 0.9$ admitted

**for** $c_k \in C$ **do**
$scoreMax \leftarrow \max_{r_i \in c_k}(score(r_i))$;
**end**
**while** $\mu(c_k) > score^{ext}_{clus}$ **do**
$newLayer_{extension} \leftarrow \{\}$;
**for** $r_j \in \{neighbors\ of\ c_k\}$ **do**
**if** $score^{ext}_{res} < score(r_j) < scoreMax$ **then**
add $r_j$ to $newLayer_{extension}$;
**end**
**end**
add $newLayer_{extension}$ to $c_k$;
$scoreMax \leftarrow \max_{r_i \in newLayer_{extension}}(score(r_i))$;
**end**
**while** $r_i \in C / r_i \in surface < 0.7 f^{DNA}_{intfrac}(x)$ **do**
iteratively filter $c_k \in C$, where $size(c_k) < size(c_{k+1})$
**end**
**if** $r_i \in C, CV_{local}(r_i) > 0.9 / r_i \in C > threshold_{buried}$ **then**
remove all residues $r_i, CV_{local}(r_i) > 0.9$ from $R$ and restart clustering;
**end**
merge clusters < 5Åaway from each other;

**Extension of a cluster $c_k$**

$score(r_i)$, score of the residue $r_i$

$\mu(c_k)$, mean score computed over a cluster $c_k$

$score^{ext}_{res}$, threshold score for residues during extension step

$score^{ext}_{clus}$, threshold score for clusters during extension step

$\{neighbors\ of\ c_k\}$, ensemble of residues < 5Å away from $c_k$

$f^{DNA}_{intfrac}(x)$, expected size of the interface given $x$ surface residues

$outLayer \leftarrow \{\}$;
**for** $c_k \in C$ **do**
**for** $r_j \in \{neighbors\ of\ c_k\}$ **do**
**if** $\frac{\mu(c_k) \times |c_k| + score(r)}{|c_k| + 1} \geq \mu(c_k)$ **then**
add $r_j$ to $outLayer$;
**end**
**end**
add $outLayer$ to $c_k$;
**end**
**if** $r_i \in C / r_i \in surface < 0.7 f^{DNA}_{intfrac}(x)$ **then**
relax $score^{seed}_{res}$, $score^{seed}_{clus}$, and $score^{ext}_{clus}$;
restart clustering;
**end**

**Addition of an outer layer to a cluster $c_k$**

$\{neighbors\ of\ c_k\}$, ensemble of residues < 5Å away from $c_k$

Table S3: **Confidence levels with which $score_{res}$ and $score_{clus}$ are determined in each layer.** The thresholds are in terms of $f^{DNA}_{intfrac}(x)$, that represents the expected size of a protein-DNA interface for a protein with $x$ surface residues. Dashes mean that the threshold cannot be relaxed.

| | Layer | First stage (not relaxed) | Second stage (relaxed) |
|---|---|---|---|
| $score_{res}$ | *seed* | $f^{DNA}_{intfrac}(x)$ | $2f^{DNA}_{intfrac}(x)$ |
| | *extension* | $2f^{DNA}_{intfrac}(x)$ | – |
| | *outer layer* | $2f^{DNA}_{intfrac}(x)$ | – |
| $score_{clus}$ | *seed* | $f^{DNA}_{intfrac}(x)/6$ | $f^{DNA}_{intfrac}(x)/4$ |
| | *extension* | $f^{DNA}_{intfrac}(x)/4$ | $f^{DNA}_{intfrac}(x)/3$ |
| | *outer layer* | $f^{DNA}_{intfrac}(x)/4$ | $f^{DNA}_{intfrac}(x)/3$ |

Table S4: **$\mathbf{JET^2_{DNA}}$ automated clustering procedure and completion by a second scoring scheme.**

**First round to detect main clusters**

```
choose D-SC1;
detect cluster seeds;
if μ(T_JET^Seeds) < 0.3 then
    choose D-SC3;
    detect cluster seeds;
end
if chosen D-SC1 then
    if μ(CV_glob^Seeds) > 0.6 and μ(PC_DNA^Seeds) < 0.9 then
        choose D-SC2;
        detect cluster seeds;
    end
end
add the extensions to the seeds;
add the outer layers to the clusters;
```

• Test whether the evolutionary signal of detected seeds is too low:

$\mu(T^{Seeds}_{JET})$, mean value of evolutionary conservation over all seeds;

• Test whether seeds are located in a concave region of the protein surface and they do not have a very high average $\mathrm{PC}_{DNA}$ value:

$\mu(PC^{Seeds}_{DNA})$, mean value of propensities over all seeds;

$\mu(CV^{Seeds}_{glob})$, mean value of global circular variance computed with $r_c = 100$Å, over all seeds;

**Second round to detect additional clusters using a complementary D-SC**

```
if chosen D-SC1 or chosen D-SC2 then
    choose D-SC3;
else
    choose D-SC1;
end
detect cluster seeds;
add the extension to the seeds;
add the outer layers to the clusters;
combine clusters;
```

Merge main clusters with additional clusters $< 5$Å away

Table S5: **Comparison of iJET$^2_{\mathrm{DNA}}$, multiVORFFIP, DISPLAR and DRNApred performances on HR-PDNA187()(*)(**)(***) and APO82 datasets.** Statistical performance values are given in percentages. To fairly assess DISPLAR, multiVORFFIP and DRNApred performances, the proteins used for training these methods were removed from HR-PDNA187. We used a sequence identity cutoff of 95% and ended up with 106 (*), 87 (**) and 42 (***) proteins, respectively. iJET$^2_{\mathrm{DNA}}$ predictions were obtained from a consensus of 2 or 8 runs out of 10. The three scoring schemes were systematically used and the best patch or combination of patches was retained. For DISPLAR and DRNApred, binary prediction results were evaluated. For multiVORFFIP, predicted residues were defined as the ones with probability > 0.5. For each dataset, the best values are highlighted in bold. If multiple values in the same column of the same dataset are reported in bold, it means that their difference, if any, is not statistical significant (p-value ≥ 0.05).

| | **Sens** | **PPV** | **Spe** | **Acc** | **F1** | **MCC** |
|---|---|---|---|---|---|---|
| **HR-PDNA187** | | | | | | |
| iJET$^2_{\mathrm{DNA}}$(2/10) | **69** | 58 | 86 | 84 | **61** | **52** |
| iJET$^2_{\mathrm{DNA}}$(8/10) | 63 | **63** | **90** | **85** | **61** | **52** |
| **HR-PDNA187*** | | | | | | |
| iJET$^2_{\mathrm{DNA}}$(2/10) | **70** | 57 | 85 | 84 | **62** | **52** |
| iJET$^2_{\mathrm{DNA}}$(8/10) | 64 | **62** | 89 | **85** | **61** | **52** |
| DISPLAR | 50 | **62** | **92** | **85** | 52 | 46 |
| **HR-PDNA187**** | | | | | | |
| iJET$^2_{\mathrm{DNA}}$(2/10) | **71** | 55 | 84 | 83 | **60** | **50** |
| iJET$^2_{\mathrm{DNA}}$(8/10) | 63 | 58 | 88 | **84** | **59** | **49** |
| multiVORFFIP ($p > 0.5$) | 44 | **65** | **93** | **84** | 50 | 43 |
| **HR-PDNA187***** | | | | | | |
| iJET$^2_{\mathrm{DNA}}$(2/10) | **70** | 54 | 82 | 81 | **60** | **48** |
| iJET$^2_{\mathrm{DNA}}$(8/10) | 62 | **59** | **87** | **82** | **59** | **48** |
| DRNApred | 36 | 46 | **86** | 78 | 25 | 35 |
| **APO82** | | | | | | |
| iJET$^2_{\mathrm{DNA}}$(2/10) | **69** | 54 | 88 | 86 | **58** | **52** |
| iJET$^2_{\mathrm{DNA}}$(8/10) | 63 | 58 | 92 | **88** | **59** | **52** |
| DISPLAR | 41 | 54 | 94 | *87* | 43 | 39 |
| multiVORFFIP ($p > 0.5$) | 44 | **64** | **95** | **87** | 50 | 45 |
| DRNApred | 23 | 40 | **95** | 85 | 24 | 21 |

Table S6: **iJET$^2_{\mathbf{DNA}}$ performance on bound and unbound forms having the same or a different stoichiometry.** HOLO82$^+$ and APO82$^+$: 74 structures having the same stoichiometry in the HOLO and APO forms. HOLO82$^{++}$ and APO82$^{++}$: 8 structures having a different stoichiometry in the HOLO and APO forms. Statistical performance are given in percentages. The performance values are reported for the default (iJET$^2_{\text{DNA}}$), the automated (iJET$^2_{\text{DNA}\,Auto}$), the complete (iJET$^2_{\text{DNA}\,Complete}$) and the automated+complete (iJET$^2_{\text{DNA}\,AutoComplete}$) clustering procedure of the algorithm. Predictions were obtained from a consensus of 2, 5 or 8 runs out of 10. The three scoring schemes were systematically used and the best patch or best combination of patches was retained.

| | **Consensus (/10 runs)** | **Sens** | **PPV** | **Spe** | **Acc** | **F1** | **MCC** |
|---|---|---|---|---|---|---|---|
| **HOLO82$^+$ (74)** | | | | | | | |
| | 2 | 67 | 55 | 90 | 87 | **59** | 52 |
| iJET$^2_{\text{DNA}}$ | 5 | 64 | 58 | 91 | 87 | **59** | 53 |
| | 8 | 61 | **60** | 93 | **88** | **59** | 53 |
| **APO82$^+$ (74)** | | | | | | | |
| | 2 | 69 | 54 | 89 | 87 | 58 | 52 |
| iJET$^2_{\text{DNA}}$ | 5 | 66 | 55 | 90 | 87 | **59** | 52 |
| | 8 | 63 | **57** | 92 | **88** | **59** | 52 |
| **HOLO82$^{++}$ (8)** | | | | | | | |
| | 2 | 70 | 55 | 82 | 79 | **59** | 47 |
| iJET$^2_{\text{DNA}}$ | 5 | 63 | 58 | 86 | **81** | **59** | 47 |
| | 8 | 60 | **62** | 89 | **81** | **59** | 48 |
| **APO82$^{++}$ (8)** | | | | | | | |
| | 2 | 73 | 52 | 83 | 82 | 59 | 50 |
| iJET$^2_{\text{DNA}}$ | 5 | 68 | 57 | 88 | 84 | **61** | 52 |
| | 8 | 64 | **59** | **90** | **84** | **60** | 51 |

Table S7: **Variation in iJET$^2_{\text{DNA}}$ performance when progressively adding layers in the clustering procedure, on bound and unbound forms.** Statistical performance values are reported for HR-PDNA187 and APO82 datasets and are given in percentages. iJET$^2_{\text{DNA}}$ predictions were obtained from a consensus of 2, 5 or 8 runs out of 10. The three scoring schemes were systematically used and the best patch or best combination of patches was retained. The performance values obtained when running predictions with an increasing number of layers in the clustering procedure are reported (1=*seed*; 2=*seed+extension*; 3 (default, also referred as iJET$^2_{\text{DNA}}$ in the other tables)=*seed+extension+outer layer*).

| | **# layers** | **Consensus (/10 runs)** | **Sens** | **PPV** | **Spe** | **Acc** | **F1** | **MCC** |
|---|---|---|---|---|---|---|---|---|
| **HR-PDNA187** | | | | | | | | |
| iJET$^2_{\text{DNA}}$ | 1 | 2 | 33 | 66 | 96 | 84 | 42 | 38 |
| | | 5 | 28 | 69 | 97 | 84 | 38 | 36 |
| | | 8 | 25 | **71** | **98** | 83 | 35 | 34 |
| | 2 | 2 | 46 | 63 | 93 | 84 | 51 | 43 |
| | | 5 | 41 | 65 | 95 | 84 | 48 | 42 |
| | | 8 | 37 | 67 | 96 | 84 | 45 | 41 |
| | 3 | 2 | **69** | 58 | 86 | 84 | **61** | **52** |
| | | 5 | 66 | 61 | 88 | **85** | **61** | **52** |
| | | 8 | 63 | 63 | 90 | **85** | **61** | **52** |
| **APO82** | | | | | | | | |
| iJET$^2_{\text{DNA}}$ | 1 | 2 | 30 | 60 | 97 | 87 | 38 | 36 |
| | | 5 | 27 | 64 | **98** | 87 | 36 | 35 |
| | | 8 | 24 | **67** | **98** | 87 | 33 | 34 |
| | 2 | 2 | 44 | 56 | 94 | 87 | 47 | 41 |
| | | 5 | 40 | 61 | 96 | 87 | 46 | 41 |
| | | 8 | 37 | 63 | 96 | 87 | 44 | 40 |
| | 3 | 2 | **69** | 54 | **88** | 86 | 58 | **52** |
| | | 5 | 66 | 55 | 90 | 87 | **59** | **52** |
| | | 8 | 63 | 58 | 92 | **88** | **59** | **52** |

Table S8: **Comparison of iJET$^2_{DNA}$, multiVORFFIP, DISPLAR and DRNApred performances on TEST-NABP82, TEST-DBP49, TEST-RBP33 and TEST-DBP24 datasets.** Statistical performance values are given in percentages. To fairly assess DISPLAR, multiVORFFIP and DRNApred performances, TEST-DBP24 and TEST-RBP33 are non redundant at 25% to our HR-PDNA187 dataset. Moreover, all the TEST sets are evaluated on a different definiton of experimental interface, defined in [3], than the one used in HR-PDNA187. iJET$^2_{DNA}$ predictions were obtained from a consensus of 2 or 8 runs out of 10. The three scoring schemes were systematically used and the best patch or combination of patches was retained. DRNApred and multiVORFFIP were evaluated on the results provided by their RNA-specific algorithms for the RNA-binding proteins, and on the results provided by their DNA-specific algorithms for the DNA-binding proteins. iJET$^2_{DNA}$ and DISPLAR, which do not have a specific algorithm to predict RNA-binding sites, were evaluated on the same algorithm for both DNA- and RNA-binding proteins. For DISPLAR and DRNApred, binary prediction results were evaluated. For multiVORFFIP, predicted residues were defined as the ones with probability $> 0.5$. For each dataset, the best values are highlighted in bold. If multiple values in the same column of the same dataset are reported in bold, it means that their difference, if any, is not statistical significant (p-value $\geq 0.05$).

| | **Sens** | **PPV** | **Spe** | **Acc** | **F1** | **MCC** |
|---|---|---|---|---|---|---|
| **TEST-NABP82** | | | | | | |
| iJET$^2_{DNA}$(2/10) | **67** | 37 | 84 | 83 | **45** | **39** |
| iJET$^2_{DNA}$(8/10) | 59 | **42** | 89 | **86** | **45** | **40** |
| DISPLAR | 37 | 33 | **91** | **86** | 30 | 25 |
| multiVORFFIP ($p > 0.5$) | 46 | 37 | 88 | **84** | 37 | 31 |
| DRNApred | 23 | 20 | **92** | **85** | 16 | 13 |
| **TEST-DBP49** | | | | | | |
| iJET$^2_{DNA}$(2/10) | **66** | 38 | 84 | 82 | **45** | **39** |
| iJET$^2_{DNA}$(8/10) | 59 | **43** | 88 | **85** | **46** | **41** |
| DISPLAR | 46 | 34 | 89 | **85** | 34 | 29 |
| multiVORFFIP ($p > 0.5$) | 49 | **42** | 89 | **85** | 41 | **35** |
| DRNApred | 30 | 28 | **92** | **85** | 23 | 19 |
| **TEST-RBP33** | | | | | | |
| iJET$^2_{DNA}$(2/10) | **68** | **36** | 84 | 83 | **44** | **39** |
| iJET$^2_{DNA}$(8/10) | 58 | **40** | 89 | **86** | **44** | **39** |
| DISPLAR | 23 | 31 | **94** | **87** | 23 | 19 |
| multiVORFFIP ($p > 0.5$) | 42 | 29 | 86 | 82 | 31 | 24 |
| DRNApred | 12 | 8 | **92** | **84** | 6 | 3 |
| **TEST-DBP24** | | | | | | |
| iJET$^2_{DNA}$(2/10) | **64** | 33 | 84 | 83 | 41 | **36** |
| iJET$^2_{DNA}$(8/10) | 57 | **41** | 90 | **87** | **44** | **39** |
| DISPLAR | 50 | **36** | **89** | **85** | 35 | **31** |
| multiVORFFIP ($p > 0.5$) | **53** | **40** | 89 | **86** | **42** | **37** |
| DRNApred | 25 | 22 | **94** | **86** | 16 | 14 |

Table S9: **Variation in iJET$^2_{\text{DNA}}$ performance when applying the default, the automated, the complete and the automated+complete clustering procedures, on bound and unbound forms.** Statistical performance values are reported for HR-PDNA187, HOLO82 and APO82 datasets and are given in percentages. The performance values are reported for the default (iJET$^2_{\text{DNA}}$), the automated (iJET$^2_{\text{DNA}\,Auto}$), the complete (iJET$^2_{\text{DNA}\,Complete}$) and the automated+complete (iJET$^2_{\text{DNA}\,AutoComplete}$) clustering procedure of the algorithm. Predictions were obtained from a consensus of 2, 5 or 8 runs out of 10. The three scoring schemes were systematically used and the best patch or best combination of patches was retained.

| | **Consensus (/10 runs)** | **Sens** | **PPV** | **Spe** | **Acc** | **F1** | **MCC** |
|---|---|---|---|---|---|---|---|
| **HR-PDNA187** | | | | | | | |
| iJET$^2_{\text{DNA}}$ | 2 | 69 | 58 | 86 | 84 | **61** | **52** |
| | 5 | 66 | 61 | 88 | **85** | **61** | **52** |
| | 8 | 63 | **63** | 90 | **85** | **61** | **52** |
| iJET$^2_{\text{DNA}\,Auto}$ | 2 | 59 | 53 | 87 | 82 | 54 | 44 |
| | 5 | 53 | 58 | 90 | 83 | 53 | 44 |
| | 8 | 46 | 62 | **93** | 84 | 50 | 43 |
| iJET$^2_{\text{DNA}\,Complete}$ | 2 | **72** | 50 | 81 | 80 | 57 | 46 |
| | 5 | 68 | 53 | 84 | 82 | 57 | 47 |
| | 8 | 64 | 54 | 86 | 82 | 57 | 47 |
| iJET$^2_{\text{DNA}\,AutoComplete}$ | 2 | 64 | 46 | 81 | 79 | 52 | 40 |
| | 5 | 58 | 49 | 85 | 80 | 51 | 40 |
| | 8 | 51 | 51 | 88 | 81 | 49 | 39 |
| **HOLO82** | | | | | | | |
| iJET$^2_{\text{DNA}}$ | 2 | 67 | 55 | 89 | 86 | **59** | **52** |
| | 5 | 65 | 58 | 91 | **87** | **59** | **52** |
| | 8 | 62 | **61** | 92 | **87** | **59** | **52** |
| iJET$^2_{\text{DNA}\,Auto}$ | 2 | 58 | 50 | 89 | 84 | 51 | 44 |
| | 5 | 52 | 55 | 92 | 86 | 51 | 44 |
| | 8 | 46 | 59 | **94** | 86 | 49 | 43 |
| iJET$^2_{\text{DNA}\,Complete}$ | 2 | **70** | 46 | 83 | 82 | 53 | 45 |
| | 5 | 67 | 48 | 85 | 83 | 54 | 48 |
| | 8 | 63 | 51 | 89 | 85 | 55 | 47 |
| iJET$^2_{\text{DNA}\,AutoComplete}$ | 2 | 62 | 43 | 84 | 81 | 49 | 39 |
| | 5 | 57 | 45 | 86 | 82 | 48 | 39 |
| | 8 | 52 | 47 | 89 | 83 | 48 | 39 |
| **APO82** | | | | | | | |
| iJET$^2_{\text{DNA}}$ | 2 | 69 | 54 | 88 | 86 | 58 | **52** |
| | 5 | 66 | 55 | 90 | 87 | **59** | **52** |
| | 8 | 63 | **58** | 92 | **88** | **59** | **52** |
| iJET$^2_{\text{DNA}\,Auto}$ | 2 | 61 | 49 | 89 | 85 | 52 | 44 |
| | 5 | 54 | 52 | 92 | 86 | 51 | 44 |
| | 8 | 47 | 55 | **94** | 87 | 49 | 43 |
| iJET$^2_{\text{DNA}\,Complete}$ | 2 | **72** | 44 | 83 | 82 | 53 | 45 |
| | 5 | 68 | 47 | 86 | 83 | 54 | 46 |
| | 8 | 64 | 49 | 88 | 85 | 54 | 47 |
| iJET$^2_{\text{DNA}\,AutoComplete}$ | 2 | 67 | 40 | 82 | 80 | 49 | 40 |
| | 5 | 60 | 43 | 86 | 82 | 48 | 39 |
| | 8 | 54 | 45 | 88 | 83 | 47 | 39 |

Table S10: **F1 values of iJET$^2_{DNA}$, multiVORFFIP (MV), DISPLAR and DRNApred for proteins presenting multiple DNA-binding sites.** F1 values are given in percentage. iJET$^2_{DNA}$ predictions were obtained from a consensus of 2 or 5 runs out of 10, depending on the protein analyzed. The three scoring schemes were systematically used and performances for each of them as well as for the best patch or best combination of patches (B.C.P.) are given. For this latter case, the scoring schemes giving the best combination of patches are reported in square brackets. The performance values obtained when running the complete clustering procedure of the program are also given in round brackets, if the corresponding F1 value is higher than the default clustering procedure. For each case studied, the first row corresponds to the DNA-binding site occupied in the structure comprised in our HR-PDNA187 dataset, the second row corresponds to the DNA-binding site(s) occupied in the alternative structure, and the third row corresponds to the union of all DNA-binding residues in the two structures. For T7 RNA polymerase having a hybrid DNA+RNA binding site, we considered DNA and RNA-binding predictions for DRNApred and multiVORFFIP since they have DNA- and RNA-specific algorithms. White lines correspond to the performance related to the DNA-binding site(s) occupied in the structure reported on top of the same column. Grey lines correspond to the performance related to the mapped DNA-binding site(s) from the alternative structure in the column beside, and to the union of all DNA-binding residues from both the structures. For DISPLAR, multiVORFFIP (MV) and DRNApred, values in red mean that the tool was trained on that complex or on a homologous one showing a similar DNA-binding site. Bold values are the best ones excluding the red ones.

| | **cplx in HR-PDNA187** | | | | | | | **alternative cplx** | | | | | | |
|---|---|---|---|---|---|---|---|---|---|---|---|---|---|---|
| | **D-SC1** | **D-SC2** | **D-SC3** | **B.C.P.** | **MV** | **DISPLAR** | **DRNApred** | **D-SC1** | **D-SC2** | **D-SC3** | **B.C.P.** | **MV** | **DISPLAR** | **DRNApred** |
| | RNA polymerase from bacteriophage T7 | | | | | | | | | | | | | |
| | **1CEZ (consensus 2/10 runs)** | | | | | | | **1MSW (consensus 2/10 runs)** | | | | | | |
| **Site 1** | 9 | 8 | 36 | 36 [D-SC3] | 81 | **39** | 3 | 8 | 11 | 37 | **41** [D-SC2+3] | 35 | 25 | 3 |
| **Site 2** | 41 (46) | 40 | 30 (0.52) | **55** [D-SC1+3] | 33 | 37 | 8 | 43 (46) | 42 (43) | 19 (49) | **50** [D-SC1+3] | 74 | 54 | 7 |
| **Union all** | 31 (36) | 31 | 31 (47) | **49** [D-SC1+3] | 60 | 44 | 5 | 34 (35) | 35 (36) | 26 (47) | **48** [D-SC1+2+3] | 69 | 48 | 5 |
| | Nuclease Domain of Adeno-Associated Virus Rep | | | | | | | | | | | | | |
| | **1UUT (consensus 5/10 runs)** | | | | | | | **1RZ9 (consensus 5/10 runs)** | | | | | | |
| **Site 1** | 44 | 9 (10) | 30 (38) | **45** [D-SC1+3] | 87 | 0 | 0 | 12 (14) | 9 | 33 | 33 [D-SC3] | **44** | 6 | 0 |
| **Site 2** | **86** | 4 (15) | 62 (84) | **86** [D-SC1] | 0 | 73 | 36 | 54 | 8 (35) | **73** | **73** [D-SC3] | 63 | 67 | 36 |
| **Union all** | **73** | 10 (20) | 49 (68) | **73** [D-SC1] | 49 | 56 | 23 | 52 | 13 (37) | 58 | 58 [D-SC3] | **78** | 54 | 23 |
| | Type 2 Restriction Endonuclease DpnI | | | | | | | | | | | | | |
| | **4ESJ (consensus 2/10 runs)** | | | | | | | **4KYW (consensus 2/10 runs)** | | | | | | |
| **Site 1** | 38 | 16 (24) | 31 (33) | **40** [D-SC1+3] | 32 | 34 | 19 | **52** | 3 (15) | 43 (**52**) | **52** [D-SC1] | 31 | 5 | 19 |
| **Site 1+2** | 64 | 50 (57) | 34 (61) | **67** [D-SC1+3] | 51 | 33 | 28 | 57 (65) | 44 (62) | 47 (**69**) | 66 [D-SC1+2+3] | 53 | 16 | 27 |
| **Union all** | 63 | 50 (58) | 36 (62) | **68** [D-SC1+3] | 55 | 38 | 29 | 55 (63) | 45 (62) | 45 (**67**) | 65 [D-SC1+2+3] | 54 | 17 | 28 |
| | Cyclic GMP-AMP synthase | | | | | | | | | | | | | |
| | **4K98 (consensus 5/10 runs)** | | | | | | | **4LEY (consensus 5/10 runs)** | | | | | | |
| **Site 1** | 0 (6) | 15 (17) | 20 | 22 [D-SC2+3] | 10 | **35** | 16 | 6 (7) | 24 | 18 | 29 [D-SC2+3] | 28 | **30** | 15 |
| **Site 1+2** | 27 | 10 | **36** | **36** [D-SC3] | 35 | 28 | 22 | 18 (20) | 40 | 29 | 44 [D-SC2+3] | **50** | 24 | 22 |
| **Union all** | 25 | 12 (14) | **37** | **37** [D-SC3] | 32 | 26 | 21 | 17 (20) | 42 | 27 | 45 [D-SC2+3] | **47** | 26 | 20 |

Table S11: Comparison of HR-PDNA187 and HOLO-APO82 with the most used datasets in previous studies.

| Datasets | #(cplx) | Resol.(Å) | Seq. Id.(%) | #(APO forms) | #(shared cplx with HR-PDNA187) | #(shared APO forms with APO82) | #(flawed cplx)* | #(cplx after PISCES filtering)*** |
|---|---|---|---|---|---|---|---|---|
| HR-PDNA187 | 187 | 2.5 | 25 | 86 | - | - | - | |
| PDNA62 [4, 5] | 62 | 3.0 | 25 | 0 | 0 | 5/62 | 2/62 | 27 |
| DBP374 [6] | 374** | 3.5 | 25 | 0 | 0 | 36/374 | 6/374 | 121 |
| MetaDBsite316 [7] | 316** | 3.0 | 30 | 0 | 0 | 53/316 | 7/316 | 139 |
| Displar264 (chains) [8] | 264 | NA | 50 | 0 | 0 | 52/264 | 12/264 | 124 |
| PDDB1.2 [9] | 47 | 3.0 | 60 | 47 | 8/47 | 8/47 | 0/47 | 28 |
| DBP206 [10] | 206** | 3.0 | 25 | 83 | 85/206 | 6/83 | 9/206 | 141 |
| PDNA224 [11] | 224** | 3.0 | 25 | 0 | 0 | 46/224 | 7/224 | 128 |
| DNABINDPROT54 [12] | 54** | NA | NA | 54 | 9/54 | 10/54 | 1/54 | 30 |

* : complexes present in the analyzed dataset, but removed from HR-PDNA187 because the asymmetric unit does not contain at least one biological unit.
**: number of non redundant chains, instead of complexes.
***: number of complexes after PISCES filtering fixing the same parameters used to filter HR-PDNA187
